# Evenness and Taylor’s law scaling shape biodiversity-stability relationships in subtropical estuarine communities

**DOI:** 10.64898/2026.05.21.726860

**Authors:** Hannah L. Bleth, Masami Fujiwara, Mark Fisher, Hui Liu, Fernando Martinez-Andrade, Joshuah S. Perkin

## Abstract

Understanding the mechanisms that link biodiversity to ecological stability is crucial for predicting ecosystem responses to global change. Using three decades of standardized fish and invertebrate monitoring data from eight subtropical estuaries in Texas, we analyze diversity components (richness, evenness, dissimilarity, and variance-mean scaling) and interpret stability patterns through synchrony or portfolio mechanisms. Regional diversity (γ-diversity) increased steadily over time, reflecting sustained gains in estuarine assemblages. Community stability, defined here as community invariability, was strongly and consistently predicted by Shannon diversity index but not by species richness, consistent with the stabilizing role of evenness. Portfolio effects were consistently strong, with community stability averaging approximately threefold higher than mean population stability, and the strength of this effect more than doubled with each unit increase in Shannon diversity. Structural equation models paired with a null model showed that Shannon diversity reduced species synchrony less than expected from statistical averaging alone, consistent with shared environmental forcing, yet its direct effect on community stability exceeded the null expectation. Taylor’s law scaling showed that abundant, persistent taxa such as blue crab (*Callinectes sapidus*), brown shrimp (*Farfantepenaeus aztecus*), and pinfish *(Lagodon rhomboides*) exhibited higher baseline invariability, contributing to community buffering, while rarer, more variable taxa introduced volatility. In contrast, compositional turnover strongly eroded stability, with high Bray-Curtis dissimilarity predicting reductions in community stability. Together, these results show that long-term estuarine community stability emerges from the combined effects of portfolio averaging, demographic variance scaling of dominant species, and the persistence of community composition, suggesting the central role of evenness-weighted diversity in biodiversity-stability relationships.

## Introduction

Ecological stability, specifically defined in this study as community invariability rather than resistance or resilience, is an important ecological property (Tilman, 1996; Tilman, 1999) linked to the long-term sustainability of ecosystems (Holling, 1973; Folke et al., 2004; Donohue et al., 2013). In coastal areas, ecological stability is especially important as these systems provide nursery habitats for numerous species (e.g., Beck et al., 2001; Minello et al., 2003; Minello et al., 2012), including ecologically and commercially important species, and buffer shorelines against physical disturbances (e.g., Barbier et al., 2011; Arkema et al., 2013). Coastal habitats are affected by both land and sea (Sheaves, 2009), and their spatiotemporal heterogeneity promotes high biodiversity. However, understanding the mechanisms that enhance or erode ecological stability in these and other dynamic environments remains a major challenge for ecology (McCann, 2000).

One key determinant of ecological stability is species biodiversity (Elton, 1927; MacArthur, 1955; Isbell et al., 2011; Cardinale et al., 2012). The classical insurance effect (Yachi and Loreau, 1999) hypothesizes that diverse communities are more stable because species respond differently to fluctuating environments. As some species decline, others increase, offsetting each other and maintaining overall community stability. This compensation results from differential responses of species to spatiotemporally heterogeneous environments, and it is expressed as a reduced species synchrony, the degree to which species abundances rise and fall together through time.

An alternative to the insurance effect, the portfolio effect, may also enhance community stability (Doak et al., 1998; Tilman, 1999; Koellner and Schmitz, 2006; Schindler et al., 2010). Under the portfolio effect, stability is produced by statistical averaging of population fluctuations. This effect can arise even when species exhibit similar responses to environmental variation, as statistical averaging reduces the influence of any single population on the community. This process is enhanced when species composition is evenly distributed, preventing a single volatile species from dominating the aggregate statistics.

Distinguishing between the insurance and portfolio effects is essential for understanding how biodiversity influences ecological stability, as diversity-stability associations can arise from statistical averaging alone or from compensatory dynamics with distinct ecological processes (Doak et al., 1998; Tilman et al., 1998). The insurance effect is strongly influenced by the loss of species with unique responses or functional roles, whereas the portfolio effect can remain insensitive to such species losses but may be more sensitive to uneven species distributions. However, both effects often use the same underlying variance and covariance structure among species abundance, so the indices for the insurance and portfolio effects are not fully statistically independent. In particular, low species synchrony can arise under both effects, either because species fluctuate independently of one another or because declines in some species are offset by increases in others. Low synchrony alone therefore cannot identify which one is operating. Consequently, we attempt to compare observed covariance patterns with a null expectation to test for effects beyond statistical averaging.

Another process associated with community stability is compositional turnover, the degree to which species’ relative abundances change over time (Diamond, 1969; Connell and Sousa, 1983; Dornelas et al., 2014; Gotelli et al., 2017). High compositional turnover can be associated with stable communities under the insurance effect because functionally similar species may replace each other. Conversely, high compositional turnover can indicate unstable communities if species fluctuate synchronously and disturbances repeatedly reset community composition. Empirical evidence of the dissimilarity-stability relationship is mixed, and few studies have examined it in long-term, large-scale datasets, especially in estuarine systems.

A third factor determining community stability operates at the level of individual populations rather than through interactions among species. Taylor’s law describes an empirical pattern in which the variance of population abundance scales as a power function of its mean (Taylor, 1961), with the exponent describing how fast variance grows with abundance. Because community-level invariability is estimated from the combined abundances of all species, taxa that are both abundant and stable (relative to their abundance) can more strongly influence the aggregate, while rarer taxa contribute less to total abundance but can still introduce instability. Taylor’s law scaling, therefore, provides a demographic basis to understand why some species contribute to community-level stability more or less than others, independently of any compensatory or averaging processes, offering a complementary view to the insurance and portfolio effects.

Recent syntheses have emphasized that biodiversity-stability relationships can be decoupled across spatial scales and between aggregate and compositional measures of variability. Wisnoski et al. (2023) tested these ideas by jointly analyzing aggregate and compositional variability across local and regional scales using long-term data of multiple taxa and ecosystem types. They found that diversity stabilized aggregate properties more than compositional ones, with regional stability depending more on spatial synchrony than diversity itself. Our study builds on this framework by applying a similar statistical approach to multi-decadal monitoring data from Texas estuaries, while extending it in three ways. First, we incorporate the Shannon diversity index (hereafter Shannon diversity), in addition to species richness, to capture species evenness, evaluating how relative abundances shape portfolio and insurance effects. Second, we use Bray-Curtis dissimilarity (Bray and Curtis, 1957) to investigate how compositional turnover influences long-term stability. Third, we empirically estimate Taylor’s law coefficients in estuarine populations (Taylor, 1961; Cohen, 2020) to test its population-level contribution to community-level stability. Together, these extensions allow us to test whether biodiversity-stability relationships identified by Wisnoski et al. (2023) across desert, forest, and deep marine ecosystems also hold in estuarine systems, while providing new insights into how evenness, compositional change, and demographic variance scaling contribute to stability under high environmental variability and disturbance.

We use long-term, large-scale monitoring data from the major bays along the Texas coast (Martinez-Andrade, 2018). These subtropical bays span a broad latitudinal gradient and experience multiple forms of disturbance. These include chronic (press) disturbances such as warming, salinity changes, reduced dissolved oxygen, and sea-level rise; acute (pulse) disturbances, such as hurricanes, deep freezes, and hypoxia; and human-induced disturbances, including fishing and pollution. Previous studies reported gradual increases in fish and invertebrate diversity due to the northward expansion of warmer-water species and contraction of colder-water species (Fujiwara et al., 2019). Consequently, the bays are undergoing large compositional changes in fish and invertebrates, though not uniformly across bays or seasons. The dynamic nature of this system provides a natural experiment-like setting for examining biodiversity-stability relationships.

Here, we address three objectives: (1) quantify the biodiversity-stability relationship, testing for effects beyond statistical averaging, and evaluating the added role of Shannon diversity; (2) evaluate how Bray-Curtis dissimilarity is associated with stability across seasons and bays; and (3) examine the demographic basis of stability by testing Taylor’s law scaling of mean-variance relationships across populations. Because the portfolio and insurance effects are both calculated from the same abundance data, apparent biodiversity-stability relationships can arise from statistical averaging alone, even when observed patterns may point to compensatory dynamics. We therefore used a variance-covariance partitioning framework paired with a null model to compare observed community stability with that expected under randomized species covariance, evaluating whether diversity influences community stability solely through statistical averaging or also through ecological mechanisms acting on species covariance. Our work has policy implications, given the strong role long-term studies play in informing conservation and management actions (Hughes et al., 2017; Fujiwara et al., 2019).

## Methods

### Data

The analysis uses long-term fisheries-independent data collected by the Texas Parks and Wildlife Department (TPWD) Coastal Fisheries Division as part of their Marine Resource Monitoring Program (Martinez-Andrade, 2018). Sampling locations were (1) Sabine Lake, (2) Galveston Bay, (3) Matagorda Bay (excluding East Matagorda Bay), (4) San Antonio Bay, (5) Aransas Bay, (6) Corpus Christi Bay, (7) Upper Laguna Madre, and (8) Lower Laguna Madre, designated as major areas 1–8, respectively (Appendix S2: Fig. S1). These bays encompass the entire Texas coast, spanning approximately 600 km. Although intensive monitoring began in 1982, the analysis uses bag seine data from 1992 to 2024 because consistent monthly sampling began in 1992. Bag seines primarily capture small-bodied species and juveniles of larger species in shallow nearshore habitats. Previous analyses of this dataset were reported by Fujiwara et al. (2019, 2022); the present study extends those analyses by examining community stability.

Sampling followed a stratified random design, with each of the eight major bays as a stratum and 20 samples collected per month per bay: 10 during the first half (days 1–15) and 10 during the second half (days 16–31). Each bay was divided into 1-minute by 1-minute grids, randomly selected without replacement each month. Each grid was further divided into 5-second by 5-second “gridlets,” and within each selected grid, one gridlet containing ≥ 15.2 m shoreline was randomly selected for sampling. This design and temporally dispersed sampling ensured sample independence and avoided pseudoreplication.

Sampling was conducted using an 18.3 m bag seine (1.8 m depth; mesh sizes 19 mm in the wings and 13 mm in the bag) following standardized protocols described by Martinez-Andrade (2018) and Fujiwara et al. (2019, 2022). All captured individuals were identified to the lowest taxonomic level possible (often species), and abundances (number of individuals per haul) were recorded.

### Data assembly and stratification

Each haul (one net deployment) recorded species-level counts with associated year, month, and bay (major area). Sample × species matrices were constructed for each bay, treating zeros as true absences. Species observed in ≤5 hauls coastwide were excluded to minimize the influence of ultra-rare taxa.

Time (year) was stratified into 11 discrete, mutually exclusive (non-overlapping) three-year periods (P1 = 1992–1994 through P11 = 2022–2024) and four seasons: Winter (Dec–Feb), Spring (Mar–May), Summer (Jun–Aug), and Fall (Sep–Nov). Three-year periods were chosen because each stability estimate requires multiple years of data to capture year-to-year variability; a shorter window would not allow variance to be estimated. A longer window, such as four years, would reduce the number of periods available across the 33-year study to fewer than 10, which would not provide enough temporal points to test for long-term trends. Autocorrelation function (ACF) checks on model residuals indicated that temporal autocorrelation between consecutive discrete blocks was weak (Lag-1 autocorrelation ≈ 0.12), indicating that the discrete periods successfully minimized temporal dependence and removed the need for an explicit autoregressive correlation structure.

A stratum is defined as the combination of bay × period × season, yielding 352 strata. Of these, 16 contained fewer than 180 samples and were excluded from diversity and stability analyses, leaving 336 strata. Eight of the excluded strata corresponded to all eight bays in the Spring 2022–2024 period (P11), which was entirely missing due to two missing months of sampling during the COVID-19 pandemic; the remaining eight were spread across four other period × season combinations, which retained five to seven of their eight bays. Consequently, of the 44 possible period × season combinations (11 periods × 4 seasons), 43 contain data: 39 with all eight bays, while four included five to seven bays. This filtering was necessary because low sample sizes can bias stability estimates, and removing these strata ensured comparability across periods and seasons.

### Diversity, stability, and turnover indices

#### Diversity metrics

Biodiversity was quantified at two spatial scales using sample-size-standardized measures.

Within-bay (α) diversity was calculated as species richness and Shannon diversity (Shannon entropy, H′), defined as

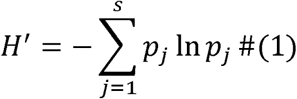

where *p_j_* is the relative abundance of species *j*, and *s* is the total number of species.

Diversity metrics were calculated from observed species abundances within each stratum. Species richness was the total number of species observed, and Shannon diversity was calculated as Shannon entropy (H′, Eq. 1) using the diversity function in the vegan package (Oksanen et al., 2025). Sampling effort was strictly controlled by the 3-year design: 20 hauls per month × 3 months per season × 3 years yields exactly 180 standardized samples per fully sampled stratum. This balanced design supports reliable variance estimation while inherently controlling for sampling effort, removing the need for rarefaction.

Regional diversity (γ-diversity) was computed by pooling across bays within each period × season without rarefaction (see Data assembly and stratification, above, for the resulting 43 period × season combinations). As a sensitivity check, we also fit the γ-diversity trend models restricted to the 39 period x season combinations in which all eight bays were sampled.

#### Stability metrics

Stability was quantified using the invariability measure of Wang et al. (2017), which expresses stability as the inverse of relative temporal variability and links naturally to spatial scaling (Wang and Loreau, 2014, 2016). Throughout this study, community stability refers to the invariability of aggregate total abundance, summed across all species within a stratum, and not to the stability of species composition or diversity. Then, compositional change is addressed separately with the Bray-Curtis dissimilarity analysis as described below. Population invariability *(I_İ_*) is determined as

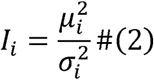

where *μ_İ_* and 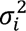 are the mean and variance of monthly total abundances of species *i* (summed across hauls) within that stratum. Therefore, stability captures variability in abundance within each bay, period, and season. Species with 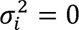 were excluded from Eq. 2 because I◻ is undefined for them.

Community invariability (*I_C_*) is determined as

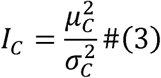

where μ*_C_* and 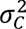 are the mean and variance of the monthly total community abundance (the number of individuals summed across species) within that stratum.

#### Portfolio effect

The portfolio effect quantifies how biodiversity buffers variability through statistical averaging (Doak et al., 1998; Tilman, 1999; Thibaut and Connolly, 2013). Here, we quantify it as the ratio of community invariability to abundance-weighted mean population invariability:

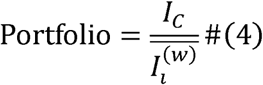

where 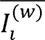 is the abundance-weighted mean of species-level invariabilities within a stratum:

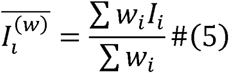

where *w_i_* = *μ_i_*. Weighting by abundance reduces the influence of rare species (often with unstable or undefined IO) and ensures the denominator of the portfolio effect reflects the dominant species. The sums in Eq. 5 are taken over species with defined IO. Portfolio > 1 indicates that averaging over species increases stability at the community level relative to the average of the constituent populations.

#### Species synchrony

Species synchrony (*φ*) was calculated following Loreau and de Mazancourt (2008), using the normalized metric *φ_LM_* bounded between 0 and 1, where φ = 1 indicates perfect synchrony (species fluctuate in the same direction at the same time) and φ = 0 indicates perfect compensation (increases in some species are exactly offset by decreases in others). For each stratum,

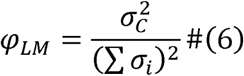

where 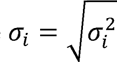 is the temporal standard deviation of species *i,* the sum in the denominator is taken over all species in a stratum, and 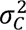 is the variance of total community abundance (Eq. 3). Species with 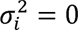, which were excluded from Eq. 2, do not affect the synchrony calculation; a species with zero temporal variance contributes zero to the sum of species standard deviations and a constant to the community total, so it changes neither the denominator nor the numerator of Eq. 6. This definition scales the variance ratio to avoid values exceeding 1.

#### Relationship between portfolio and synchrony metrics

The portfolio effect (Eq. 4) is the ratio of community-level to abundance-weighted mean population-level invariability. The ratio increases whenever species within a stratum fluctuate less than fully synchronously, regardless of the underlying mechanism producing the lack of full synchrony. Consequently, the portfolio effect alone does not distinguish two sources of stability: the purely statistical effect of averaging over multiple populations, and ecological compensatory dynamics. Because species synchrony (φ) is calculated from the same underlying variance terms as the portfolio effect (Eqs. 4-6), the two metrics are mathematically related rather than statistically independent.

This mathematical relationship is most transparent at the limiting case of a single species: with only one species present, community-level and population-level invariability are identical by definition (I_c_ = I_i(w)_), so Portfolio = 1, and species synchrony likewise reduces to φ = 1, since there is only one variance term in both the numerator and denominator of Eq. 6. As richness or evenness increases from this boundary, Portfolio can rise above 1 while φ falls from 1 toward 0, but this divergence reflects the mechanical structure of the indices as much as any underlying ecological process.

To address this, we generated null datasets that preserved species-level variances and diversity while removing true covariance among species (see Synchrony, diversity, and community stability associations), allowing us to set a baseline expectation for stability arising from statistical averaging alone. Departures of observed stability metrics from this null baseline indicate a contribution from non-random, compensatory dynamics among species, beyond what would be expected from averaging over multiple populations.

#### Log-transformation of stability and synchrony metrics

Synchrony was log-transformed on the natural logarithmic scale (log□ φ), and community stability on the base-10 scale (log□□ I_C_). Synchrony and community stability were logged on different bases. This has no consequence for inference, because a change of base is a linear rescaling that affects only the units of the coefficients. The log transformation was motivated by the structure of the metrics; because *σ_C_*^2^ = φ(Σσ□)², community invariability satisfies log I_C_ = 2 log μ_C_ − log φ − 2 log(Σσ□) for any consistent base, so log φ enters community stability linearly by construction. The log transformation also reduces the pronounced right skew of φ. Although φ is bounded on [0, 1], observed values ranged from 0.037 to 0.976 across the 336 strata, the upper bound also retains its interpretation on the transformed scale, since log φ = 0 corresponds to perfect synchrony. A floor of ε = 1×10□□ was applied before logging to guard against undefined values at φ = 0, but no stratum required it. The portfolio effect and population invariability were similarly log□□ -transformed for the same variance-stabilizing purpose in the analyses described below.

#### Temporal turnover (Bray-Curtis Dissimilarity)

Temporal β-diversity was quantified as Bray-Curtis dissimilarity (Bray and Curtis, 1957) between consecutive years on a monthly scale. For each bay × month, species abundances were pooled across hauls, and absent species were assigned zero abundance. Dissimilarities were calculated only between consecutive years within the same bay × month combination and then averaged across months within each bay × period × season (stratum) to yield a single dissimilarity estimate per stratum. Bray-Curtis dissimilarity was computed as

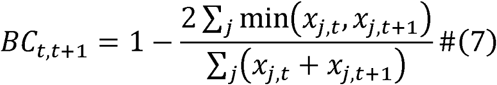

where *x_j,t_* is the abundance of species *j* at year *t*, using the “vegdist” function in the vegan package. Restricting to adjacent years emphasizes short-term dissimilarity and reduces confounding by long-term directional trends, ensuring comparability with the stability metric anchored to the same temporal scale. This approach does not capture compositional change that takes place gradually over longer time spans, such as shifts associated with press disturbances. Evaluating longer-term trajectories is left for future work. For comparison, we also calculated dissimilarity by pooling data across all months within each bay × season, but results were qualitatively unchanged.

### Modeling and inference

All statistical analyses were conducted in R (R Core Team, 2024). ChatGPT-5 (OpenAI) was used interactively to assist with debugging and annotating R code during data analysis. Then, Claude Opus 5 (Anthropic) was used to review and revise the R code. All code and results were checked by the authors.

### Regional γ-diversity trends over time

Temporal trends in regional (γ) diversity were evaluated using richness and Shannon metrics pooled across bays within each season and 3-year period. The 3-year grouping was chosen to match the temporal window used for estimating invariability and to reduce within-season and interannual fluctuations. To capture nonlinear long-term patterns, we fitted generalized additive models (GAMs) using the “gam” function in the mgcv package (Wood, 2017). Each model used γ-diversity as the response and mid-year of each 3-year period as the predictor, representing time, to test whether γ-diversity changed over the 33-year study period, with a thin-plate regression spline smoother (k=6) and restricted maximum likelihood (REML) estimation. Separate GAMs were fitted for each season (n = 11 periods per season, except n = 10 for Spring, because period P10 was excluded; see Study design). Because γ-diversity was pooled across bays, no random terms were included. Linear regressions were also fitted to estimate overall slopes, and false-discovery-rate (FDR) corrections (Benjamini and Hochberg 1995) were applied across seasons, with q-values reported as FDR-adjusted p-values.

### Objective 1: Biodiversity-stability relationships (portfolio and insurance effects)

#### Diversity vs. community stability

We evaluated how local diversity relates to community stability by linking α-diversity to community invariability (*I_C_*). We fitted separate season-specific linear mixed-effect models for each diversity metric (richness and Shannon diversity), using community stability as the response and diversity as the predictor. Each model included bay as a random effect (season-specific n ranged from 80 to 88, reflecting the retained strata per season; see Table 1). Eight models were fitted in total (two diversity metrics × four seasons), and false-discovery-rate corrections were applied, with q-values reported as FDR-adjusted p-values.

**Table 1.**
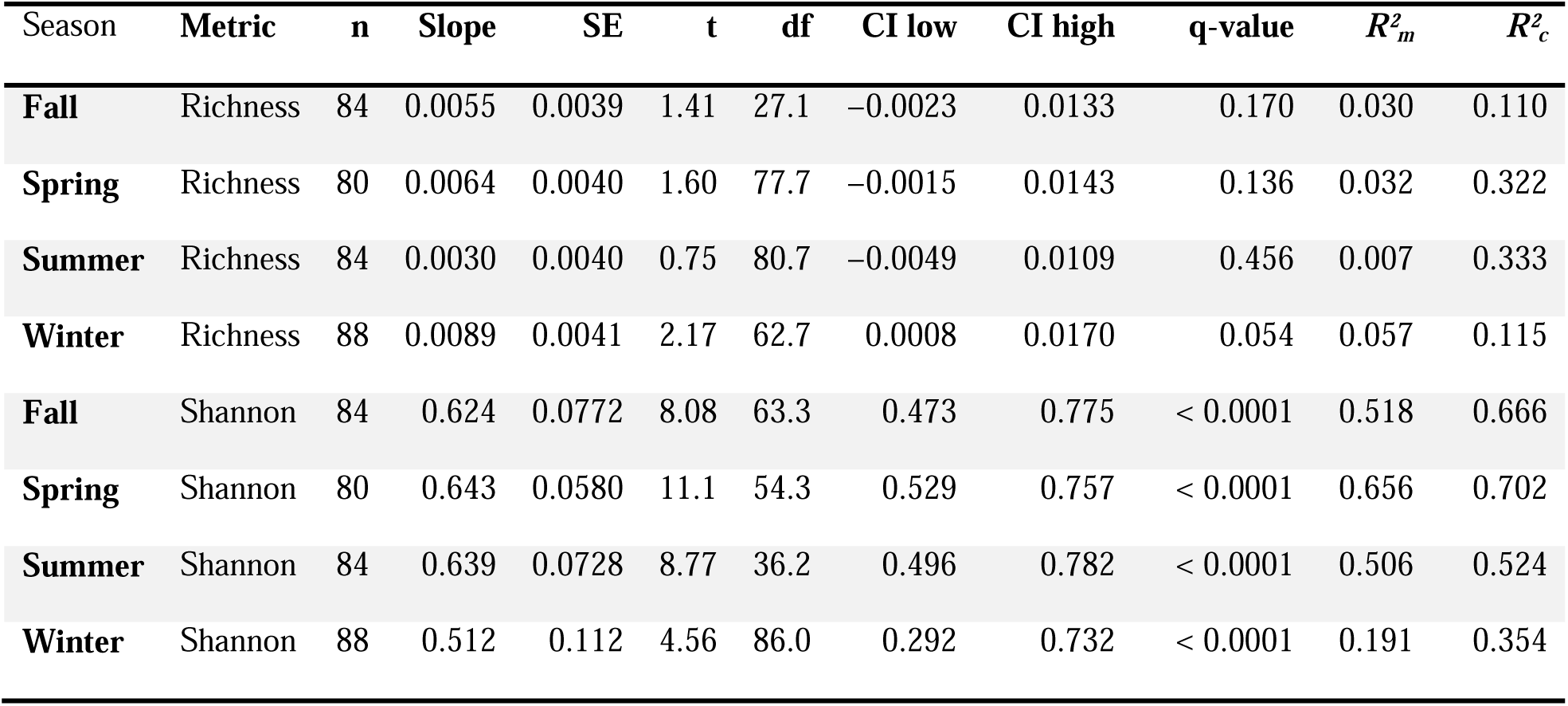
Season-specific diversity-stability slopes from linear mixed-effects models. Response variable: log_10_ *I_C_* (community invariability). Predictors: richness or Shannon diversity. Models: log_10_ *I_C_* ∼ predictor + (1 | major_area), fitted separately by season. Slopes are fixed-effect estimates with Satterthwaite df; 95% confidence intervals are Wald (estimate ± t-critical × SE). Reported significance is based on q-values (Benjamini-Hochberg adjusted across the eight seasonal tests). Marginal R² (*R²*□) denotes the proportion of variance explained by fixed effects (diversity alone), whereas conditional R² (*R²*\*) represents the variance explained by both fixed and random effects (diversity + among-bay differences).

#### Temporal trend in portfolio effect

We evaluated whether the portfolio effect changed through time using a generalized additive model (GAM). The response was the log_10_-transformed portfolio effect, and predictors included a smooth term for mid-year of each three-year period and categorical effects for season and bay (major area). The smooth term used a thin-plate regression spline (k = 6), fitted using REML in the mgcv package (Wood, 2017). Season and bay were included to account for systematic seasonal and spatial differences in stability. For comparison, a generalized least-squares (GLS) model with a linear term for time, season, and bay was also fitted to test for an overall linear trend.

#### Portfolio effect vs. diversity

To assess whether diversity underlies these dynamics, we fit mixed-effects models relating the log_10_ portfolio effect to richness and Shannon diversity. Three models were evaluated: one with richness, one with Shannon diversity, and one including both predictors. Because Shannon diversity incorporates both richness and evenness, some correlation between Shannon diversity and richness is expected. Including both in the same model tests whether Shannon diversity predicts the portfolio effect beyond what is attributable to richness alone, which points toward a contribution of evenness, although this analysis does not isolate evenness as an independent predictor from richness. Each model included random intercepts for bay and season to account for repeated measures across space and time. The analysis included 336 observations (see Table 1 for the breakdown by season), fitted using the “lmer” function of the lme4 package (Bates et al., 2015).

#### Synchrony, diversity, and community stability associations

To quantify diversity-synchrony associations, we modeled log(φ) as a function of diversity using linear mixed-effects regressions. Two models were fit with log(φ) as the response and either richness or Shannon diversity as the fixed effect, with random intercepts for bay and season, using REML in the nlme package (Pinheiro et al., 2025). Predictors were left on their original scales for interpretability.

We evaluated whether diversity influenced stability directly and/or indirectly via synchrony using two piecewise structural equation models (SEMs): one with richness and one with Shannon diversity. Each SEM comprised two mixed-effects submodels with random intercepts for bay and season. Similar to the log(φ) models, season was modeled as nested within bay (32 groups) in these SEM submodels, rather than crossed with bay as in the diversity-stability and portfolio models. Standardized path coefficients (STD β) were obtained by standardizing variables prior to coefficient extraction, enabling comparison across paths. The analysis used the piecewiseSEM package (Lefcheck, 2016). To isolate ecological mechanisms from statistical coupling, we generated 1,000 randomized datasets. Within each stratum, the monthly abundance time series of each species was permuted independently across months.

Because within-species permutations reorder the same set of monthly values, it preserves each species’ mean and temporal variance as well as both Shannon diversity and the denominator of Eq. 6, while removing any covariance among species. Community invariability and synchrony were recalculated from the randomized series using Eqs. 3 and 6, and standardized path coefficients were obtained from piecewise SEMs using the same specification as for the observed data, establishing a baseline of pure statistical averaging. One-tailed permutation p-values were calculated as (1 + r)/(1 + n), where r is the number of null replicates at least as extreme as the observed coefficient.

### Objective 2: Compositional turnover and stability

#### Turnover (Bray-Curtis Dissimilarity)-stability association

We tested whether temporal dissimilarity constrains community stability using linear mixed-effects models. Turnover was measured as mean Bray-Curtis dissimilarity between years *t* and *t*+1, computed at the bay (major area) × season × 3-year period level. Stability was measured as community invariability, *I_c_*, for the corresponding period. We fit a linear mixed-effects model with log_10_,(*I_C_*) as the response with dissimilarity as the predictor, season as a fixed effect to account for seasonal differences, and a random intercept for bay to absorb spatial baselines. This model evaluates whether higher dissimilarity predicts lower stability while controlling for seasonal variation and spatial heterogeneity among bays.

Although both metrics derive from the same underlying monthly abundance data, their variance structures are not identical. Community invariability’s variance term (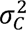, Eq. 3) pools variation across all nine monthly totals within a stratum, combining both within-season (month-to-month) and between-year variation without separating them. Bray-Curtis dissimilarity, in contrast, isolates the between-year axis specifically, comparing the same calendar month across consecutive years and excluding within-season comparisons entirely. Year-to-year variation therefore contributes to, but does not solely determine, 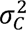, so the two measures share underlying data without being mathematically identical.

### Objective 3: Demographic basis of stability (Taylor’s law scaling)

#### Abundance-variability scaling

We tested whether species with larger populations were more stable, following Taylor’s law (Taylor, 1961). Population variance (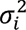) was regressed against mean abundance (*μ_i_*) on a log_10_-log_10_ scale, excluding zero values. *μ_i_* and 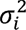 are the monthly mean and variance of species *i* abundance defined in Eq. 2. The Taylor’s law exponent *b* was estimated from 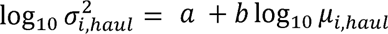. Because 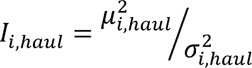, log invariability scales with log mean abundance with slope *m* = 2 − *b*, so b < 2 (m > 0) means more abundant populations are relatively more stable. We interpreted slopes in light of theoretical developments linking stochastic population dynamics to Taylor’s law (Cohen et al., 2013) and more recent syntheses (Cohen, 2020).

## Results

A total of 301 taxa was recorded across multiple broad biological categories (Appendix S1: Table S1). The majority were bony fishes (174 taxa), with 1 identified only to class, 15 to family, 4 to genus, and 154 to species. Crustaceans were also well represented (62 taxa), followed by mollusks (47 taxa) and cnidarians (7 taxa). Smaller contributions came from cartilaginous fishes (5 species), tunicates (2 species), and single representatives of annelid worms, echinoderms, insects, and reptiles.

### Regional γ-diversity trends over time

Regional γ-diversity showed clear long-term increases across Texas bays, with both richness and Shannon diversity contributing to this pattern. GAMs revealed significant nonlinear temporal structure in γ-diversity that varied among seasons (Appendix S2: Fig. S2; Appendix S1: Table S2). For richness, smooth terms were significant across all seasons (edf = 1.0–2.4, ref.df = 1.0–2.9, F = 7.8–18.7, p-values ranging from < 0.0001 to 0.0016; adjusted R² = 0.84, deviance explained = 87.6%, n = 43). For Shannon diversity, smooth terms were also significant (edf = 2.6–4.2, ref.df = 3.2–4.7, F = 5.6–8.8, p-values ranging from < 0.0001 to 0.0039; adjusted R² = 0.77, deviance explained = 85.3%, n = 43). These results indicate nonlinear temporal variation in γ-diversity superimposed on long-term trends.

Linear regressions provided complementary evidence of sustained increases across seasons. Richness increased at 1.21 species · year□¹ (95% Wald CI: [0.66, 1.76], t□□ = 4.27, p < 0.0001, R² = 0.31, F□,□□ = 18.25), while Shannon diversity increased by 0.0177 per year (95% Wald CI: [0.0083, 0.0271], t□□ = 3.71, p <0.001, R² = 0.25, FO□,□□ = 13.7). Restricting this analysis to the 39 complete combinations gave a similar richness slope (1.35 vs. 1.21 species ·year□¹, both p < 0.001), indicating that the varying bay coverage did not affect the conclusion. Together, these results demonstrate that regional diversity increased consistently over three decades, with nonlinear, season-dependent fluctuations captured by the GAM smooths.

### Objective 1: Biodiversity-stability relationships (portfolio and insurance effects)

#### Diversity vs. community stability

Community stability (log10,(*I_C_*)) increased with diversity, but only Shannon diversity consistently and strongly predicted stability across all seasons (Fig. 1; Table 1). For richness, the diversity-stability relationship was positive but weak, with slopes ranging from 0.003 to 0.009 and generally non-significant after multiple-comparison correction. The winter model showed the steepest slope (β = 0.0089, 95% CI: [0.0008, 0.0170], t□□.□ = 2.17, q = 0.054), suggesting a marginal seasonal effect, whereas slopes for fall, spring, and summer were small and non-significant. In contrast, Shannon diversity showed steep, highly significant positive slopes in every season (0.51–0.64, all 95% CIs within [0.29, 0.78], all q < 0.0001), indicating that communities with greater evenness-weighted diversity were more stable. Marginal R² (0.19–0.66) shows that Shannon diversity alone explained a large proportion of variance in stability, while conditional R² (0.35–0.70) demonstrates that additional variation was captured by differences among bays.

**Figure 1.**
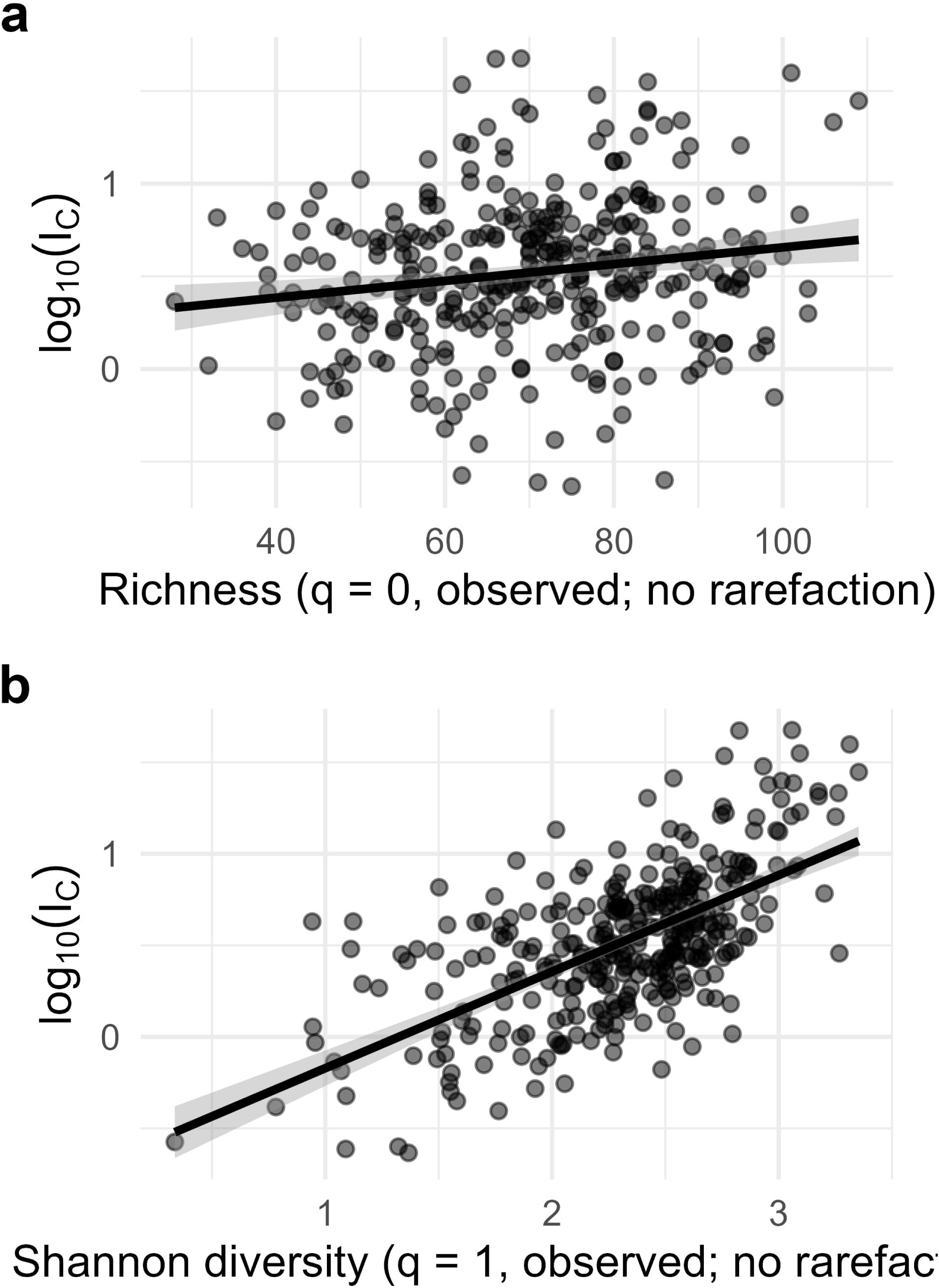
Diversity-stability relationships (pooled across seasons). Points represent bay × period × season observations; line is the OLS fit with 95% confidence interval. (a) Community stability (log_10_ *I_C_*) versus species richness. (b) Community stability (log_10_ *I_C_*) versus Shannon diversity.

#### Temporal trend in portfolio effect

Across bays and seasons, the portfolio effect consistently exceeded 1, indicating persistent statistical averaging. A GAM on log□□(Portfolio) showed significant nonlinear temporal structure (smooth for mid-year: F□.□□, □.□□ = 4.82, p < 0.001), explaining 27.4% of deviance (adjusted R² = 0.242; n = 336; Appendix S2: Fig. S3; Appendix S1: Table S3). Seasonal and spatial differences were evident: relative to Fall, portfolio values were lower in Spring (β = −0.141, 95% CI: [−0.215, −0.067], t□□□ = −3.73, p = 0.0002) and Summer (β = −0.110, 95% CI: [−0.183, −0.037], t□□□ = −2.94, p = 0.0035), but not Winter (β = −0.046, 95% CI: [−0.118, 0.026], t□□□ = −1.26, p = 0.21). A complementary linear model indicated a modest overall increase embedded within these complex, nonlinear multi-decadal fluctuations (β = 0.00426 log□□ units yr□¹, 95% CI: [0.0014, 0.0071], t□□□ = 2.96, p = 0.003), equivalent to ∼1.0% yr□¹ on the original scale (10^0.00426^ ≈ 1.010). Across all 336 strata, the mean portfolio effect was 3.22 (SE = 0.125; median = 2.65, bootstrap 95% CI: [2.42, 2.81]) and significantly exceeded 1 (one-sided Wilcoxon signed-rank test, W = 55,930, p < 0.0001), suggesting a strong portfolio effect across the study system.

#### Portfolio effect vs. diversity

Linear mixed models relating log□□(Portfolio effect) to diversity (n = 336; random intercepts for bay and season) showed contrasting patterns (Fig. 2). With richness as the sole predictor (Fig. 2a), the effect was weak (β = 0.0022, 95% CI: [−0.0004, 0.0049], t_108.9_ =1.67, p = 0.099; R²m = 0.015, R²c = 0.235), implying that adding one species increases the portfolio effect by only ∼0.5% (10^0.00223^ ≈ 1.005). The Shannon model (Fig. 2b) showed a strong positive association (β = 0.314, 95% CI: [0.260, 0.368], t_76.0_ = 11.40, p < 0.0001, R²m = 0.304, R²c = 0.351), indicating that a one-unit increase in Shannon roughly doubles the portfolio effect (10^0.314^ ≈ 2.06). In the combined model, Shannon remained strongly positive (β = 0.338, 95% CI: [0.284, 0.392], t_42.7_ = 12.27, p < 0.0001; R²m = 0.321, R²c = 0.353), whereas richness was slightly negative and non-significant (β = −0.0019, 95% CI: [−0.0040, 0.0002], t_37.5_ = −1.78, p = 0.084). These results indicate that evenness-weighted diversity, rather than species counts, is the primary correlate of a stronger portfolio effect (Appendix S1: Table S4).

**Figure 2.**
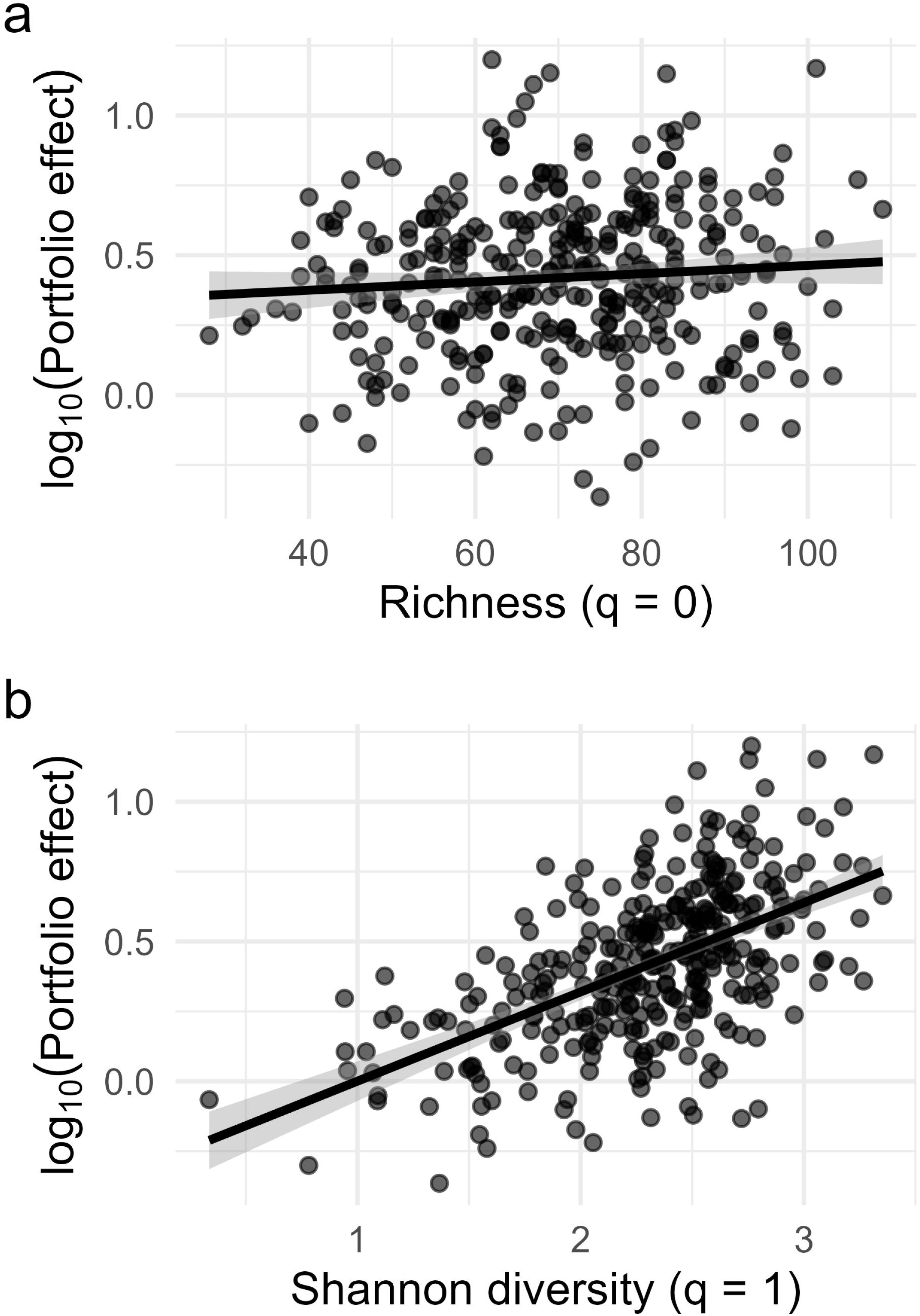
Portfolio effect vs. diversity (n = 336). Points are bay × period × season observations; lines show marginal linear mixed-effects fits with 95% confidence intervals (calculated from fixed effects only, excluding random effect variance). (a) log_10_ (Portfolio) versus species richness. (b) log_10_ (Portfolio) versus Shannon diversity.

#### Synchrony, diversity, and community stability associations

Linear mixed-effects regressions revealed that synchrony declined with increasing diversity (n = 336; Fig. 3). Synchrony varied among strata but remained inside its theoretical bounds, ranging from 0.037 to 0.976 (mean = 0.278, median = 0.238), with only one stratum above 0.95 and five below 0.05. For richness, the relationship was weak but significant (β = −0.0072, 95% CI: [−0.0125, −0.0019], t□□□ = −2.66, p = 0.008; R²□ = 0.03, R²c = 0.32), indicating only a slight decline in log(φ) with increasing richness (Fig. 3a). Shannon diversity showed a strong negative relationship with synchrony (β = −0.902, 95% CI: [−1.014, −0.790], t□□□ = −15.7, p < 0.0001; R²□ = 0.48, R²c = 0.55), indicating that communities with higher evenness-weighted diversity fluctuated markedly less synchronously (Fig. 3b). These results indicate that evenness-weighted diversity explains a greater proportion of variation in synchrony than richness, consistent with a stronger role of evenness in promoting asynchrony among species.

**Figure 3.**
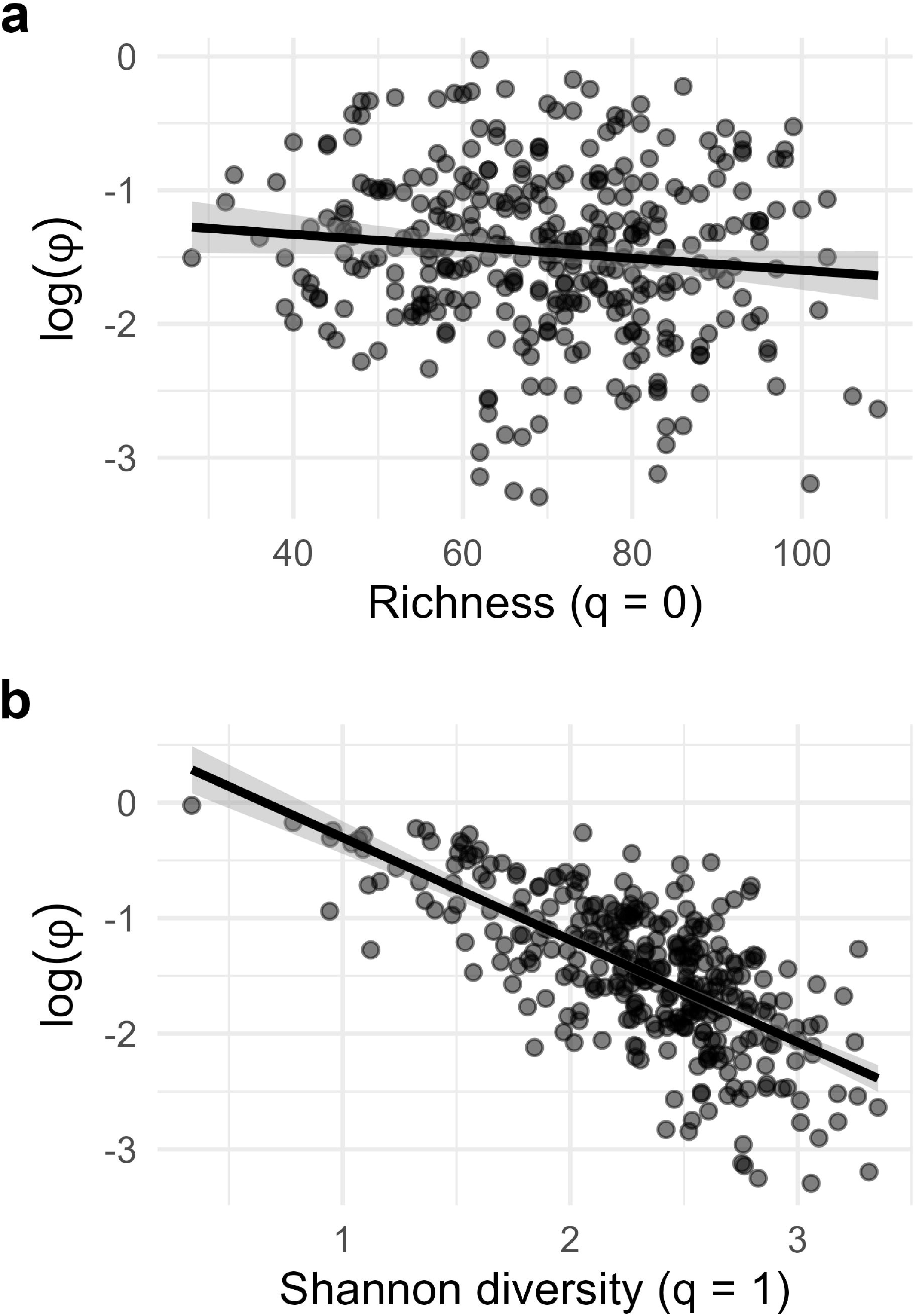
Diversity-synchrony relationships. Points are bay × period × season observations; lines show marginal linear mixed-effects fits with 95% confidence intervals (calculated from fixed effects only, excluding random effect variance). (a) log_e_ φ versus species richness. (b) log_e_ φ vs Shannon diversity.

We evaluated whether diversity influences stability directly and/or indirectly through synchrony using piecewise SEM with log-transformed synchrony (log φ) and log-transformed community invariability (log_10_ (*I_C_*)). Models were fit separately with diversity measured as richness or Shannon diversity (n = 336), accounting for bay and season (Appendix S1: Table S5; Fig. 4). In both models, synchrony exerted a strong negative effect on stability (i.e., greater asynchrony among species was stabilizing). However, the pathways through which diversity influenced stability differed between richness and Shannon diversity.

**Figure 4.**
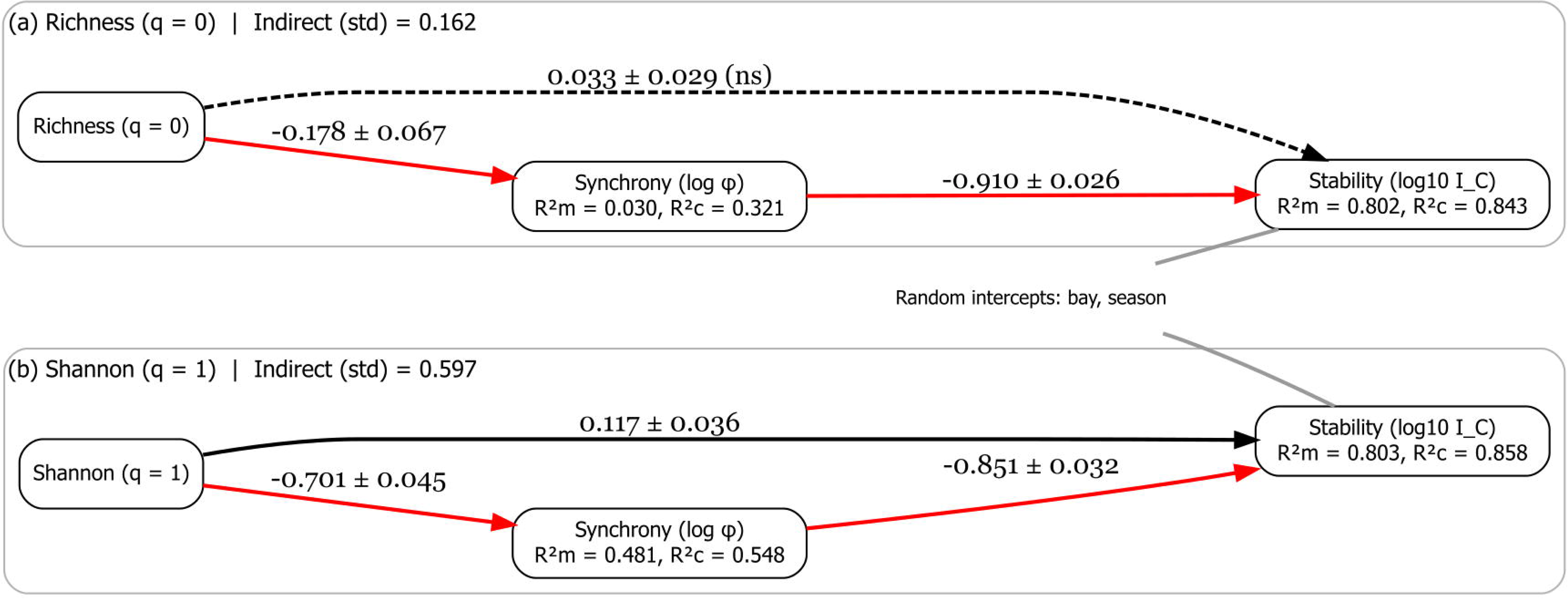
Piecewise structural equation model (SEM) path diagrams. Arrows show standardized path coefficients (STD β); dashed lines indicate non-significant paths. Indirect effect is the product of the diversity → synchrony and synchrony → stability coefficients. Random intercepts for bay and season; node labels show marginal and conditional R².

In the richness SEM, richness had no detectable direct effect on stability (STD β = +0.033, 95% CI: [−0.024, 0.090], t_302_ = 1.02, p = 0.31). Richness was associated with a weak but significant reduction in synchrony (STD β = −0.178, 95% CI: [−0.309, −0.047], t_303_ = −2.66, p = 0.008), and because synchrony strongly destabilized stability (STD β = −0.911, 95% CI: [−0.962, −0.860], t_302_ = −35.42, p < 0.0001), richness indirectly promoted stability through this pathway. The indirect effect was +0.162, the product of the standardized richness-to-synchrony and synchrony-to-stability coefficients (–0.178 × –0.911). Richness, therefore, explained little variation in synchrony (R²□ = 0.03) and had no detectable direct effect on stability, so its overall contribution was weak despite a statistically detectable mediated pathway.

The Shannon SEM revealed much stronger patterns. Shannon diversity had a significant direct positive effect on stability (STD β = +0.117, 95% CI: [0.046, 0.188], t_302_ = 3.23, p = 0.0014) and a strong negative effect on synchrony (β = −0.701, 95% CI: [−0.789, −0.613], t_303_ = −15.71, p < 0.0001). Because log-synchrony strongly reduced stability (β = −0.851, 95% CI: [−0.914, −0.788], t_302_ = −26.21, p < 0.0001), the indirect effect of Shannon diversity was substantial: +0.597, the product of the standardized Shannon-to-synchrony and synchrony-to-stability coefficients (–0.701 × –0.851). Shannon diversity was thus positively associated with stability through both direct and indirect pathways.

Because all possible pathways between the three variables were estimated, both SEMs were fully saturated (just-identified), precluding the need for a global goodness-of-fit test (e.g., Fisher’s C). However, individual submodel fits were strong: marginal R^2^ for stability was ≈0.80–0.81, while R^2^ for synchrony was ≈3% in the richness model but ≈48% in the Shannon model. These results indicate that evenness-weighted diversity is the dominant predictor of stability, acting directly and indirectly by promoting asynchrony, whereas richness explained little variation in synchrony and had no detectable direct effect.

Importantly, because synchrony depends on covariance among species, our SEM framework explicitly partitions the diversity-stability relationship into changes in species covariance (the indirect synchrony pathway) and remaining stabilizing effects (the direct pathway). Under the null model, pure statistical averaging predicted that Shannon diversity would strongly reduce synchrony (mean null STD β = –0.811, 95% null CI: [–0.849, –0.771]).

The observed reduction in synchrony (STD β = –0.701, 95% CI: [–0.789, –0.613]) was weaker than every one of the 1,000 null replicates (one-tailed permutation p = 1.000 against the hypothesis that it was stronger). This pattern is consistent with shared environmental forcing inducing positive covariance among species, which offsets part of the reduction expected under independence. In contrast, statistical averaging alone predicted essentially no direct effect of Shannon diversity on stability (mean null STD β = 0.006, 95% null CI: [–0.033, 0.043]). Our observed direct effect fell outside the entire null distribution (STD β = 0.117, 95% CI: [0.046, 0.188], permutation p < 0.001), indicating that Shannon diversity provides ecological stabilizing effects.

### Objective 2: Compositional turnover and stability

#### Turnover (Bray-Curtis Dissimilarity)-stability association

We evaluated how compositional turnover (Bray-Curtis dissimilarity between consecutive years) related to community stability (log_10_(*I_C_)*) using a linear mixed-effects model with season as a fixed effect and bay as a random intercept (n = 336). Turnover was strongly negatively associated with stability (β = −2.470, 95% CI: [−2.848, −2.092], t_325.9_ = −12.82, p < 0.0001; Fig. 5; Appendix S1: Table S6). On the original scale, a unit increase in dissimilarity reduced *I_c_* by ∼99.7% (10□²·□□□ ≈ 0.0034), and a change of 0.1 was associated with a ∼43% decline (10□□·²□□ ≈ 0.57).

**Figure 5.**
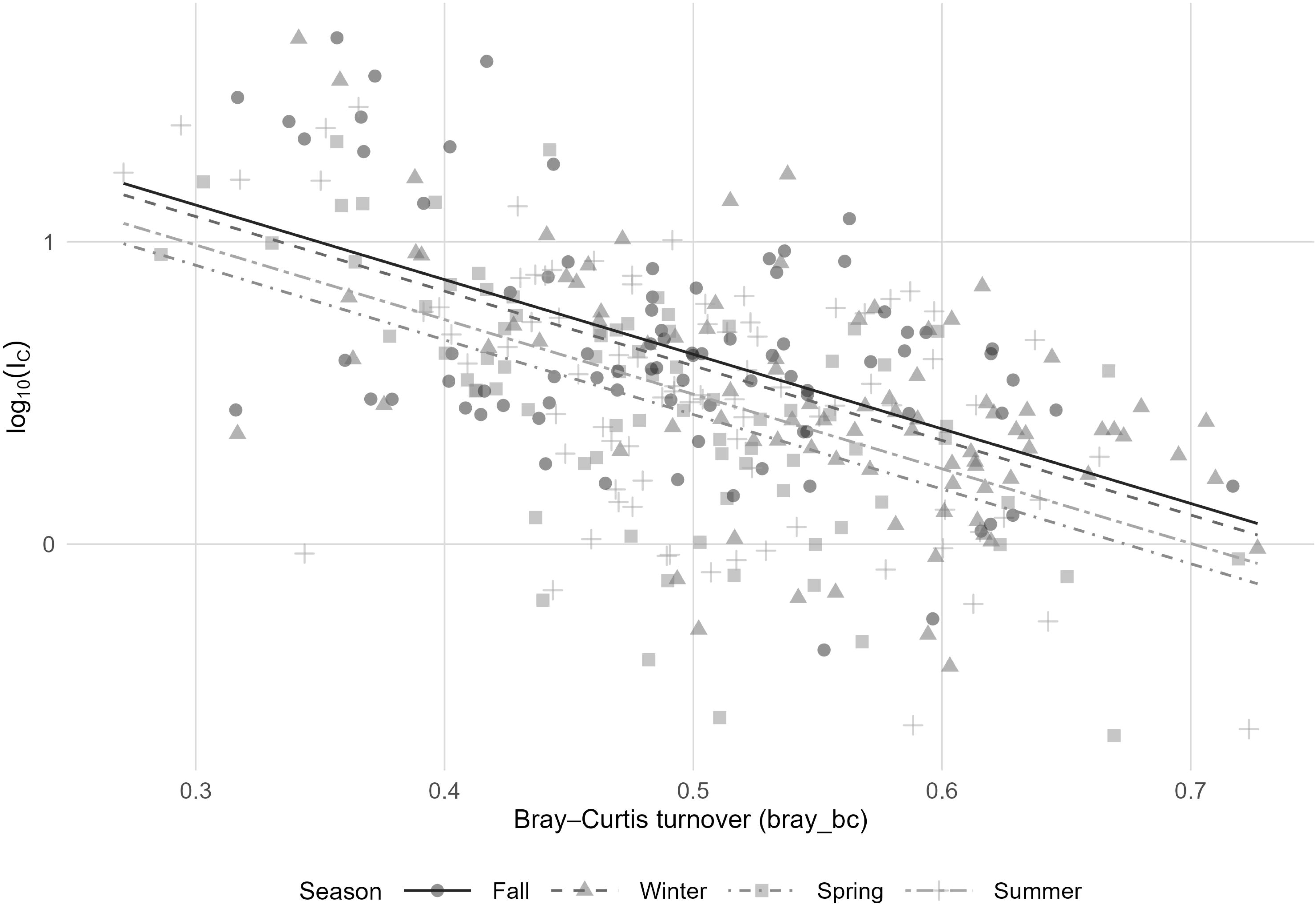
Compositional turnover vs. community stability across seasons. Points are bay × period × season observations; lines are linear mixed-effects model predictions with dissimilarity and season as fixed effects and bay as a random intercept.

Seasonal differences were evident relative to Fall: Spring showed ∼37% lower stability (β = −0.199, 95% CI: [−0.291, −0.107], t_324.0_ = −4.25, p < 0.0001), Summer ∼26% lower (β = −0.132, 95% CI: [−0.222, −0.042], t_324.1_ = −2.85, p = 0.005), and Winter did not differ significantly (β = −0.038, 95% CI: [−0.130, 0.054], t_324.1_ = −0.81, p = 0.42). Including season significantly improved model fit (Δχ²□ = 21.69, p < 0.0001). Random intercept variance among bays was 0.024 (SD = 0.156) and residual variance was 0.090 (SD = 0.300). These results show that higher year-to-year compositional turnover is strongly associated with reduced community stability, even after accounting for seasonal structure and spatial heterogeneity.

### Objective 3: Demographic basis of stability (Taylor’s law scaling)

#### Abundance-variability scaling (Taylor’s law)

We evaluated whether population variability scales with mean abundance following Taylor’s law. Across 23,494 species × bay × period × season observations, we found a strong abundance-variability relationship (Fig. 6). The Taylor’s law regression (OLS, 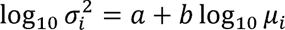) yielded b = 1.686 (95% CI: [1.682, 1.690], t_23492_ = 852.4, R² = 0.969, p < 0.0001, n = 23,494), well within the expected ecological range (1 < b < 2), indicating that variance increases predictably with the mean but less than quadratically, so more abundant species are proportionally less variable and relatively more stable. Expressed in terms of invariability 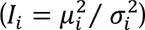, the equivalent scaling slope was m = 2 − b = 0.314, simply showing an algebraic identity rather than an independent confirmation (see Methods). Species-level contrasts further illustrated these patterns (Appendix S2: Fig. S4): abundant and persistent taxa such as blue crab (*Callinectes sapidus*), brown shrimp (*Farfantepenaeus aztecus*), and pinfish (*Lagodon rhomboides*) showed higher baseline stability, whereas less common and more variable taxa such as diamond-backed terrapin (*Malaclemys terrapin*), lane snapper (*Lutjanus synagris*), and Atlantic tripletail (*Lobotes surinamensis*) contributed to lower stability.

**Figure 6.**
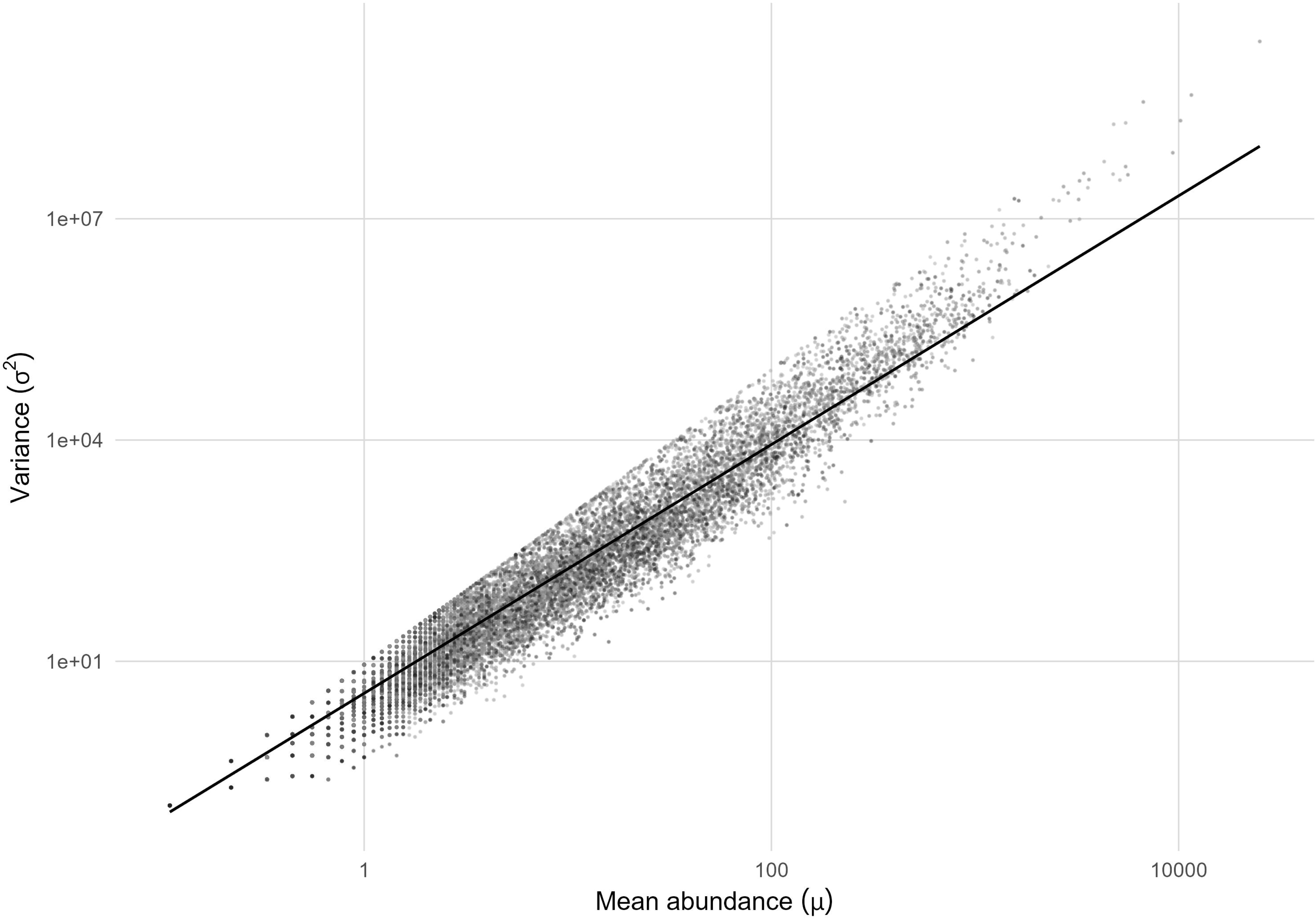
Taylor’s law scaling of variance (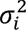) against mean abundance (*μ_İ_*) on a log-log scale. Points are species × bay × period × season observations (n = 23,494); line is the OLS fit.

## Discussion

Our analysis revealed a long-term increase in regional diversity across the subtropical estuaries, with significant increases in every season (Appendix S2: Fig. S2). This pattern is consistent with previous work showing that local diversity has been increasing in almost all bays, reflecting the increased occurrence of tropical and subtropical species and associated shifts in estuarine assemblages (Fujiwara et al., 2019; Pawluk et al., 2021). Such sustained gains provide an important basis for interpreting biodiversity-stability relationships because, as the species pool expands, the potential for portfolio averaging and compensatory dynamics may also increase.

Consistent with this expanding diversity, community stability itself was strongly and consistently predicted by Shannon diversity in every season, whereas richness showed only weak, mostly non-significant relationships (Results: Diversity vs. community stability; Table 1; Fig. 1). Marginal R² for these season-specific Shannon models ranged from 0.19 to 0.66, indicating that the strength of this relationship varied considerably across seasons even though its direction and significance were consistent. The pathways underlying this relationship are examined further below through structural equation modeling.

We found strong evidence for a portfolio effect at the community level. Community invariability was on average more than three times higher than mean population invariability, indicating that statistical averaging across species provides a stabilizing force, consistent with the portfolio hypothesis (Doak et al., 1998; Tilman, 1999; Schindler et al., 2010). The magnitude of the portfolio effect was tightly linked to Shannon diversity, whereas relationships with richness were weak and generally non-significant (Fig. 2; Appendix S1: Table S4). A one-unit increase in Shannon diversity more than doubled the portfolio effect, while richness produced no discernible increase. This pattern, together with the sign reversal of richness once Shannon is included in the same model (Results: Portfolio effect vs. diversity), suggests that the evenness component of Shannon diversity, rather than species number, is the more likely driver. When relative abundances are more balanced, species contributions are distributed more evenly across the assemblage, consistent with the stabilizing influence of averaging. This interpretation follows Jost’s (2007) broader argument that diversity indices sensitive to relative abundances, such as Shannon entropy, better capture the balance of species contributions than richness alone.

Accounting for this evenness-weighted component is useful for interpreting stability because it captures aspects of community structure, such as dominance and the distribution of relative abundances, that raw richness does not, and it also addresses concerns that richness measures are sensitive to sampling effort and dominance (Gotelli and Colwell, 2001). These results are consistent with communities having more balanced relative abundances maintaining stable aggregate abundance better under environmental fluctuations, reflecting the stronger stabilizing influence of species-level averaging associated with evenness-weighted diversity.

Piecewise SEMs further clarify the pathways by which diversity shapes stability (Appendix S1: Table S5; Fig. 4). Because species synchrony (φ) is mathematically related to the portfolio effect (Eq. 6; Methods), pure statistical averaging is expected to reduce synchrony as diversity increases, even without ecological compensation among species. Our models show that Shannon diversity reduced synchrony (STD β = −0.701), which was strongly and negatively associated with community stability (STD β = −0.851), generating a large indirect positive effect (+0.597; see null-model comparison below). Thus, Shannon diversity influences stability through both a direct positive pathway and a strong indirect pathway mediated by reduced synchrony. Richness had no direct effect and explained little variation in synchrony.

While classical theory often attributes this asynchrony to an ‘insurance effect’ driven by compensatory dynamics (Yachi and Loreau, 1999), our null model results tell a different ecological story: the observed synchrony reduction (STD β = −0.701) was in fact weaker than the −0.811 expected under pure statistical independence, indicating that shared environmental forcing, not compensatory dynamics, limits how far synchrony falls. Estuaries are harsh; many species respond in the same direction to shared drivers such as salinity, temperature, and freshwater inflow, producing positive covariance among species abundances, causing them to fluctuate somewhat together. However, evenness-weighted diversity generates such a large statistical portfolio effect that it successfully reduces overall synchrony and stabilizes the community despite this synchronizing environmental forcing.

Because diversity, community invariability, and synchrony are derived from the same abundance time series, their relationships may contain elements of statistical coupling. To address this, we used variance-covariance partitioning with a null model to establish the baseline of statistical averaging under perfect species independence. We found that statistical coupling explains part of the Shannon-based biodiversity-stability relationship, particularly the diversity-synchrony association. However, when controlling for this covariance structure, the observed direct effect of Shannon diversity on stability (STD β = 0.117) was significantly stronger than the null expectation from statistical coupling alone, which was indistinguishable from zero (STD β = 0.006, 95% null CI: [–0.033, 0.043]; p < 0.001). This indicates that the biodiversity-stability relationship is not fully explained by statistical coupling among diversity, synchrony, and stability. The source of this excess direct effect cannot be fully resolved in the current study. It may reflect a stabilizing mechanism operating independently of species synchrony. For example, communities with more balanced relative abundances may be less susceptible to any single species dominating fluctuations in total abundance. This would be a structural consequence of evenness-weighted diversity that is distinct from the synchrony pathway modeled here. Alternatively, it may reflect ecological compensation not completely captured by our synchrony metric. Distinguishing between these explanations would require experimental manipulation of species composition and evenness, which is beyond the scope of this monitoring-based study.

Taylor’s law analyses provided further demographic insight (Fig. 6). Population variance scaled predictably with mean abundance (Taylor, 1961; Cohen, 2020), with a slope less than 2 (b ≈ 1.69), indicating that common species are proportionally less variable than rare ones. This is consistent with theoretical predictions that abundant, persistent species exhibit inherently higher baseline invariability (Tilman et al., 1998). Species such as blue crab, brown shrimp, and pinfish contributed disproportionately to community buffering due to their high abundances and relatively low variance (Appendix S2: Fig. S4), while more variable and less common taxa introduced volatility.

In contrast to the stabilizing influences of evenness-weighted diversity and asynchrony, compositional turnover was negatively associated with stability (Appendix S1: Table S6; Fig. 5). Higher Bray-Curtis dissimilarity predicted lower community invariability. Part of this association could reflect shared underlying variability in the monthly abundance data, since both measures are derived from the same monthly abundance series. However, this overlap is partial rather than a direct definitional dependency, unlike the log-linear relationship between synchrony and stability described above (Methods: Log-transformation). Community invariability’s variance term pools both within-season and between-year variation, whereas Bray-Curtis dissimilarity isolates the between-year axis alone. Year-to-year compositional change is therefore one contributor to community variance among several, not its sole determinant. This association may also reflect disturbance-driven reassembly of community composition. Distinguishing the relative contributions of shared data structure versus ecological reassembly would require a null model analogous to the one used for the diversity-synchrony-stability analysis, which is left for future work. This tension between beneficial diversity effects and destabilizing compositional change is consistent with theory linking α- and β-components of diversity to stability across scales (Wang and Loreau, 2014, 2016).

Our findings extend those of Wisnoski et al. (2023), who found that diversity stabilizes aggregate community properties more than compositional ones, a relationship that can decouple across spatial scales. Similar to their study, we observed strong aggregate stability linked to diversity, while compositional turnover eroded it. Unlike the deserts, forests, and open oceans examined by Wisnoski et al. (2023), estuaries are highly dynamic and frequently exposed to natural and anthropogenic disturbances. In such volatile systems, shared environmental forcing often limits classical compensatory dynamics, making the statistical portfolio effect, with immense buffering capacity, the dominant mechanism sustaining community stability. By incorporating Shannon diversity, our study shows an additional dimension: evenness-weighted diversity is particularly important in promoting asynchrony and stability in estuarine communities. Taylor’s law analyses and taxon-specific contributions further show that abundant, persistent species provide demographic stability while more variable taxa introduce volatility. Finally, our dissimilarity analysis shows that rapid compositional change destabilizes assemblages, in contrast to the stabilizing roles of statistical averaging and compensatory dynamics.

These results extend the richness-based framework of classic theory (Tilman, 1996, 1999; Doak et al., 1998; Tilman et al., 1998; Yachi and Loreau, 1999) and modern syntheses (Loreau and de Mazancourt, 2013; Wang and Loreau, 2014, 2016; Wisnoski et al., 2023) by showing that biodiversity-stability relationships are scale- and context-dependent and sensitive to evenness-weighted diversity, demographic scaling, and temporal dissimilarity. Our analyses show that long-term stability arises not from any single mechanism but from the interplay of portfolio averaging overcoming environmental forcing, demographic contributions of dominant species, and compositional persistence. Conservation and management strategies should therefore consider not only species richness but also the evenness component reflected in Shannon diversity, asynchrony, demographic structure, and compositional persistence, as these components jointly determine long-term stability. Recognizing how these mechanisms shape variability can improve efforts to sustain the ecological functions of subtropical estuarine communities.

## Supporting Information

Additional supporting information may be found in the online version of this article:

**Appendix S1. Supplementary Tables:** Includes taxa counts by broad category, GAM outputs for regional γ-diversity and the portfolio effect, mixed-effects model summaries relating diversity to the portfolio effect, structural equation model (SEM) path coefficients, and linear mixed-effects model results for dissimilarity-stability associations.

**Appendix S2. Supplementary Figures:** Includes the study area map, temporal trends in regional γ-diversity, temporal trends in the portfolio effect (log□□ scale) with GAM smooths and confidence intervals, and species-level baseline contributions to stability.

## Acknowledgments

In accordance with institutional guidelines, MF^1^ discloses the use of ChatGPT-5 (OpenAI; accessed August-September 2025) to suggest and refine wording and sentence structure and to assist with debugging, cleaning, and annotating R code, and the use of Claude (Anthropic; accessed August 2026) during revision of the manuscript: Claude Sonnet 5 for suggestions on wording and sentence structure, and Claude Opus 5 for an independent audit of the R code, verification of reported numerical results, and corrections of R code. All AI-assisted suggestions, whether textual or related to code, were reviewed, edited, and independently verified by MF^1^ before inclusion, who assumes full responsibility for the content and accuracy of the manuscript. The manuscript was subsequently reviewed and revised by all authors. HLB was supported by the POST 9/11 GI Bill; no other funding supported this work. (*MF¹ = Masami Fujiwara)*

## Author Contributions

HLB and MF^1^ contributed equally to this project. The study was conceived by HLB and MF^1^. HLB conducted the initial analyses and drafted the manuscript as part of her M.S. thesis under the supervision of MF^1^ and committee members HL and JSP. MF^1^ subsequently refined the analyses and substantially rewrote the thesis for the manuscript. HLB, HL, and JSP made a major effort in revising the manuscript and suggesting additional analysis. MF^2^ and FMA provided the data and offered guidance on its use throughout the project.

(*Here, MF¹ = Masami Fujiwara; MF² = Mark Fisher.*)

## References

Arkema, K. K., G. Guannel, G. Verutes, S. A. Wood, A. Guerry, M. Ruckelshaus, P. Kareiva, M. Lacayo, and J. M. Silver. 2013. Coastal habitats shield people and property from sea-level rise and storms. Nature Climate Change 3:913–918. 10.1038/nclimate1944

Barbier, E. B., S. D. Hacker, C. Kennedy, E. W. Koch, A. C. Stier, and B. R. Silliman. 2011. The value of estuarine and coastal ecosystem services. Ecological Monographs 81:169–193. 10.1890/10-1510.1

Bates, D., M. Mächler, B. Bolker, and S. Walker. 2015. Fitting linear mixed-effects models using lme4. Journal of Statistical Software 67:1–48. 10.18637/jss.v067.i01

Beck, M. W., K. L. Heck Jr., K. W. Able, D. L. Childers, D. B. Eggleston, B. M. Gillanders, and B. Halpern, et al. 2001. The identification, conservation, and management of estuarine and marine nurseries for fish and invertebrates: A better understanding of the habitats that serve as nurseries for marine species and the factors that create site-specific variability in nursery quality will improve conservation and management of these areas BioScience 51:633–641. 10.1641/0006-3568(2001)051[0633:TICAMO]2.0.CO;2

Benjamini, Y., and Y. Hochberg. 1995. Controlling the false discovery rate: a practical and powerful approach to multiple testing. *Journal of the Royal Statistical Society*, Series B 57:289–300. 10.1111/j.2517-6161.1995.tb02031.x

Bray, J. R., and J. T. Curtis. 1957. An ordination of the upland forest communities of southern Wisconsin. Ecological Monographs 27:325–349. 10.2307/1942268

Cardinale, B. J., J. E. Duffy, A. Gonzalez, D. U. Hooper, C. Perrings, P. Venail, A. Narwani, et al. 2012. Biodiversity loss and its impact on humanity. Nature 486:59–67. 10.1038/nature11148

Cohen, J. E. 2020. Species-abundance distributions and Taylor’s power law of fluctuation scaling. Theoretical Ecology 13:607–614. 10.1007/s12080-020-00470-x

Cohen, J. E., M. Xu, and W. S. F. Schuster. 2013. Stochastic multiplicative population growth predicts and interprets Taylor’s power law of fluctuation scaling. Proceedings of the Royal Society B 280:20122955. 10.1098/rspb.2012.2955

Connell, J. H., and W. P. Sousa. 1983. On the evidence needed to judge ecological stability or persistence. The American Naturalist 121:789–824. 10.1086/284105

Diamond, J. M. 1969. Avifaunal equilibria and species turnover rates on the Channel Islands of California. Proceedings of the National Academy of Sciences USA 64:57–63. 10.1073/pnas.64.1.57

Doak, D. F., D. Bigger, E. K. Harding, M. A. Marvier, R. E. O’Malley, and D. Thomson.1998. The statistical inevitability of stability–diversity relationships in community ecology. The American Naturalist 151:264–276. 10.1086/286117

Donohue, I., O. L. Petchey, J. M. Montoya, A. L. Jackson, L. McNally, M. Viana, K. Healy, M. Lurgi, N. E. O’Connor, and M. C. Emmerson. 2013. On the dimensionality of ecological stability. Ecology Letters 16:421–429. 10.1111/ele.12086

Dornelas, M., N. J. Gotelli, B. McGill, H. Shimadzu, F. Moyes, C. Sievers, and A. E. Magurran. 2014. Assemblage time series reveal biodiversity change but not systematic loss. Science 344:296–299. 10.1126/science.1248484

Elton, C. S. 1927. Animal Ecology. Macmillan, London, UK.

Folke, C., S. Carpenter, B. Walker, M. Scheffer, T. Elmqvist, L. Gunderson, and C. S. Holling. 2004. Regime shifts, resilience, and biodiversity in ecosystem management. Annual Review of Ecology, Evolution, and Systematics 35:557–581. 10.1146/annurev.ecolsys.35.021103.105711

Fujiwara, M., F. Martinez-Andrade, R. J. D. Wells, M. Fisher, M. Pawluk, and M. C. Livernois. 2019. Climate-related factors cause changes in the diversity of fish and invertebrates in subtropical coast of the Gulf of Mexico. Communications Biology 2:403. 10.1038/s42003-019-0650-9

Fujiwara, M., A. Simpson, M. TorresOCeron, and F. MartinezOAndrade. 2022. LifeOhistory traits and temporal patterns in the incidence of coastal fishes experiencing tropicalization. Ecosphere 13:e4188. 10.1002/ecs2.4188

Gotelli, N. J., and R. K. Colwell. 2001. Quantifying biodiversity: procedures and pitfalls in the measurement and comparison of species richness. Ecology Letters 4:379–391. 10.1046/j.1461-0248.2001.00230.x

Gotelli, N. J., H. Shimadzu, M. Dornelas, B. McGill, F. Moyes, and A. E. Magurran. 2017. Community-level regulation of temporal trends in biodiversity. Science Advances 3: e1700315. 10.1126/sciadv.1700315

Holling, C. S. 1973. Resilience and stability of ecological systems. Annual Review of Ecology, Evolution, and Systematics 4:1–23. 10.1146/annurev.es.04.110173.000245

Hughes, B. B., R. Beas-Luna, A. K. Barner, K. Brewitt, D. R. Brumbaugh, E. B. Cerny-Chipman, S. L. Close, et al. 2017. Long-term studies contribute disproportionately to ecology and policy. BioScience 67:271–281. 10.1093/biosci/biw185

Isbell, F., V. Calcagno, A. Hector, J. Connolly, W. S. Harpole, P. B. Reich, M. Scherer-Lorenzen, et al. 2011. High plant diversity is needed to maintain ecosystem services. Nature 477:199–202. 10.1038/nature10282

Jost, L. 2007. Partitioning diversity into independent alpha and beta components. Ecology 88:2427–2439. 10.1890/06-1736.1

Koellner, T., and O. J. Schmitz. 2006. Biodiversity, ecosystem function, and investment risk. BioScience 56:977–985. 10.1641/0006-3568(2006)56[977:BEFAIR]2.0.CO;2

Lefcheck, J. S. 2016. piecewiseSEM: Piecewise structural equation modeling in R for ecology, evolution, and systematics. Methods in Ecology and Evolution 7:573–579. 10.1111/2041-210X.12512

Loreau, M., and C. de Mazancourt. 2008. Species synchrony and its drivers: Neutral and non-neutral community dynamics in fluctuating environments. The American Naturalist 172:E48–E66. 10.1086/589746

Loreau, M., and C. de Mazancourt. 2013. Biodiversity and ecosystem stability: a synthesis of underlying mechanisms. Ecology Letters 16:106–115. 10.1111/ele.12073

MacArthur, R. 1955. Fluctuations of animal populations and a measure of community stability. Ecology 36:533–536. 10.2307/1929601

Martinez-Andrade, F. 2018. Marine resource monitoring operations manual. Texas Parks and Wildlife Department, Coastal Fisheries Division, Austin, Texas, USA

McCann, K. S. 2000. The diversity-stability debate. Nature 405:228–233. 10.1038/35012234

Minello, T. J., K. W. Able, M. P. Weinstein, and C. G. Hays. 2003. Salt marshes as nurseries for nekton: testing hypotheses on density, growth, and survival through meta-analysis. Marine Ecology Progress Series 246:39–59. 10.3354/meps246039

Minello, T. J., L. P. Rozas, and R. Baker. 2012. Geographic variability in salt marsh flooding patterns may affect nursery value for fishery species. Estuaries and Coasts 35:501–514. 10.1007/s12237-011-9463-x

Oksanen, J., G. L. Simpson, F. G. Blanchet, R. Kindt, P. Legendre, P. R. Minchin, R. B. O’Hara, et al. 2025. vegan: Community Ecology Package. R package version 2.7–1, CRAN. 10.32614/CRAN.package.vegan

Pawluk, M., M. Fujiwara, and F. Martinez-Andrade. 2021. Climate effects on fish diversity in the subtropical bays of Texas. Estuarine, Coastal and Shelf Science 249:107121. 10.1016/j.ecss.2020.107121

Pinheiro, J., D. Bates, and R Core Team. 2025. nlme: Linear and Nonlinear Mixed Effects Models. R package version 3.1–168. 10.32614/CRAN.package.nlme

R Core Team. 2024. R: A Language and Environment for Statistical Computing. Vienna, Austria: R Foundation for Statistical Computing. https://www.R-project.org/

Schindler, D. E., R. Hilborn, B. Chasco, C. P. Boatright, T. P. Quinn, L. A. Rogers, and M. S. Webster. 2010. Population diversity and the portfolio effect in an exploited species. Nature 465:609–612. 10.1038/nature09060

Sheaves, M. 2009. Consequences of ecological connectivity: the coastal ecosystem mosaic. Marine Ecology Progress Series 391:107–115. 10.3354/meps08121

Taylor, L. R. 1961. Aggregation, variance and the mean. Nature 189:732–735. 10.1038/189732a0

Thibaut, L. M., and S. R. Connolly. 2013. Understanding diversity-stability relationships: towards a unified model of portfolio effects. Ecology Letters 16:140–150. 10.1111/ele.12019

Tilman, D. 1996. Biodiversity: population versus ecosystem stability. Ecology 77:350–363. 10.2307/2265614

Tilman, D. 1999. The ecological consequences of changes in biodiversity: a search for general principles. Ecology 80:1455–1474. 10.1890/0012-9658(1999)080[1455:TECOCI]2.0.CO;2

Tilman, D., C. L. Lehman, and C. E. Bristow. 1998. Diversity-stability relationships: statistical inevitability or ecological consequence? The American Naturalist 151:277–282. 10.1086/286118

Wang, S., and M. Loreau. 2014. Ecosystem stability in space: α, β and γ variability. Ecology Letters 17:891–901. 10.1111/ele.12292

Wang, S., and M. Loreau. 2016. Biodiversity and ecosystem stability across scales in metacommunities. Ecology Letters 19:510–518. 10.1111/ele.12582

Wang, S., M. Loreau, J.-F. Arnoldi, J. Fang, K. A. Rahman, S. Tao, and C. de Mazancourt. 2017. An invariability-area relationship sheds new light on the spatial scaling of ecological stability. Nature Communications 8:15211. 10.1038/ncomms15211

Wisnoski, N. I., R. Andrade, M. C. N. Castorani, C. P. Catano, A. Compagnoni, T. Lamy, N. K. Lany, et al. 2023. Diversity-stability relationships across organism groups and ecosystem types become decoupled across spatial scales. Ecology 104:e4136. 10.1002/ecy.4136

Wood, S. N. 2017. Generalized Additive Models: An Introduction with R. Second edition. Chapman & Hall/CRC, Boca Raton, Florida, USA. 10.1201/9781315370279

Yachi, S., and M. Loreau. 1999. Biodiversity and ecosystem productivity in a fluctuating environment: the insurance hypothesis. Proceedings of the National Academy of Sciences USA 96:1463–1468. 10.1073/pnas.96.4.1463

