## Appendix S1 for "Evenness and Taylor’s law scaling shape biodiversity-stability relationships in subtropical estuarine communities"

**Journal:** Ecology

**Appendix S1. Supplementary Tables**

**Table S1.** Counts of taxa by broad category and taxonomic level. The dataset includes a total of 303 taxa.

| **category** | **phylum** | **class** | **order** | **suborder/super-family** | **family** | **genus** | **species** |
| --- | --- | --- | --- | --- | --- | --- | --- |
| annelid worm | 0 | 1 | 0 | 0 | 0 | 0 | 1 |
| bony fish | 0 | 1 | 0 | 0 | 15 | 4 | 154 |
| cartilaginous fish | 0 | 0 | 0 | 0 | 0 | 0 | 5 |
| cnidarian | 0 | 1 | 1 | 0 | 0 | 0 | 5 |
| crustacean | 0 | 0 | 1 | 1 | 6 | 2 | 52 |
| echinoderm | 0 | 1 | 0 | 0 | 0 | 0 | 1 |
| insect | 0 | 0 | 0 | 0 | 0 | 1 | 0 |
| mollusk | 1 | 1 | 1 | 0 | 1 | 2 | 41 |
| reptile | 0 | 0 | 0 | 0 | 0 | 0 | 1 |
| tunicate | 0 | 0 | 0 | 0 | 0 | 0 | 2 |

**Table S2.** Summary of parametric and smooth terms from GAMs of γ-richness and γ-Shannon. Statistic = t for parametric terms and F for smooth terms. p = raw p-value; q = BH-adjusted p-value. Dashes (–) indicate terms without parametric estimates (smooth components).

| **Component** | **Term** | **Statistic** | **p-value** | **q-value** | **Estimate** | **SE** |
| --- | --- | --- | --- | --- | --- | --- |
| **Richness** | | | | | | |
| Parametric | (Intercept) | t₃₂₀ = 51.89 | < 0.0001 | < 0.0001 | 130.47 | 2.51 |
| Parametric | seasonSpring | t₃₂₀ = 8.11 | < 0.0001 | < 0.0001 | 29.63 | 3.65 |
| Parametric | seasonSummer | t₃₂₀ = 10.77 | < 0.0001 | < 0.0001 | 38.30 | 3.56 |
| Parametric | seasonFall | t₃₂₀ = 9.03 | < 0.0001 | < 0.0001 | 32.10 | 3.56 |
| Smooth | s(mid_year):seasonWinter | F₂.₃₉, ₂.₉₃ = 9.56 | < 0.001 | < 0.001 | – | – |
| Smooth | s(mid_year):seasonSpring | F₁.₄₇, ₁.₈₀ = 17.27 | < 0.0001 | < 0.0001 | – | – |
| Smooth | s(mid_year):seasonSummer | F₁.₀₀, ₁.₀₀ = 18.69 | < 0.001 | < 0.001 | – | – |
| Smooth | s(mid_year):seasonFall | F₁.₇₄, ₂.₁₅ = 7.77 | 0.0016 | 0.0016 | – | – |
| **Shannon** | | | | | | |
| Parametric | (Intercept) | t₃₂₀ = 53.19 | < 0.0001 | < 0.0001 | 2.60 | 0.0488 |
| Parametric | seasonSpring | t₃₂₀ = −3.73 | < 0.001 | 0.0025 | −0.265 | 0.0712 |
| Parametric | seasonSummer | t₃₂₀ = 2.41 | 0.023 | 0.026 | 0.167 | 0.0691 |
| Parametric | seasonFall | t₃₂₀ = 1.19 | 0.25 | 0.25 | 0.0819 | 0.0691 |
| Smooth | s(mid_year):seasonWinter | F₂.₇₉, ₃.₄₀ = 5.99 | 0.0027 | 0.0046 | – | – |
| Smooth | s(mid_year):seasonSpring | F₄.₁₈, ₄.₆₉ = 8.77 | < 0.0001 | < 0.001 | – | – |
| Smooth | s(mid_year):seasonSummer | F₂.₇₂, ₃.₃₁ = 5.62 | 0.0039 | 0.0052 | – | – |
| Smooth | s(mid_year):seasonFall | F₂.₆₁, ₃.₂₀ = 5.96 | 0.0029 | 0.0046 | – | – |

**Table S3.** GAM results for log₁₀(Portfolio effect). Generalized additive model of log₁₀(Portfolio) as a function of mid-period year (smooth term), season, and major area. Parametric terms report coefficient estimates (β), standard errors (SE), t-values, and p-values. The smooth term reports effective degrees of freedom (edf), F statistic, and p-value. Model fit: adjusted R² = 0.242; deviance explained = 27.4%; n = 336.

| **Term** | **Estimate** | **SE** | **t value** | **df** | **p-value** |
| --- | --- | --- | --- | --- | --- |
| **Parametric terms** |  |  |  |  |  |
| (Intercept) | 0.29635 | 0.04320 | 6.860 | 321 | < 0.0001 |
| seasonSpring | −0.14095 | 0.03777 | −3.732 | 321 | < 0.001 |
| seasonSummer | −0.10953 | 0.03725 | −2.941 | 321 | 0.0035 |
| seasonWinter | −0.04633 | 0.03682 | −1.258 | 321 | 0.209 |
| major_area2 | 0.01813 | 0.05235 | 0.346 | 321 | 0.729 |
| major_area3 | 0.19464 | 0.05268 | 3.694 | 321 | < 0.001 |
| major_area4 | 0.23160 | 0.05203 | 4.451 | 321 | < 0.0001 |
| major_area5 | 0.28093 | 0.05268 | 5.333 | 321 | < 0.0001 |
| major_area6 | 0.31838 | 0.05268 | 6.043 | 321 | < 0.0001 |
| major_area7 | 0.22981 | 0.05203 | 4.417 | 321 | < 0.0001 |
| major_area8 | 0.30053 | 0.05235 | 5.741 | 321 | < 0.0001 |
| **Smooth term** |  |  |  |  |  |
| s(period_mid) | edf = 3.99 | Ref.df = 4.563 | F = 4.82 |  | < 0.001 |

**Table S4.** Mixed-effects models relating log₁₀(Portfolio effect) to diversity. Linear mixed models with random intercepts for bay (major area) and season (n = 336). Fixed-effect estimates (β), standard errors (SE), denominator degrees of freedom (df), t values, and p values are shown. Marginal R² (R²m) represents variance explained by fixed effects; conditional R² (R²c) includes random effects.

| **Model** | **Predictor** | **Estimate (β)** | **SE** | **df** | **t value** | **p-value** | **R²m** | **R²c** |
| --- | --- | --- | --- | --- | --- | --- | --- | --- |
| **Richness-only model** | Intercept | 0.2627 | 0.1086 | 42.9 | 2.42 | 0.0199 | 0.015 | 0.235 |
|  | Richness | 0.00223 | 0.00134 | 108.9 | 1.67 | 0.099 |  |  |
| **Shannon-only model** | Intercept | –0.30675 | 0.07071 | 43.7 | –4.34 | < 0.0001 | 0.304 | 0.351 |
|  | Shannon | 0.31384 | 0.02753 | 76.0 | 11.40 | < 0.0001 |  |  |
| **Combined model** | Intercept | –0.23284 | 0.08345 | 23.9 | –2.79 | 0.0102 | 0.321 | 0.353 |
|  | Richness | –0.00186 | 0.00105 | 37.5 | –1.78 | 0.084 |  |  |
|  | Shannon | 0.33835 | 0.02759 | 42.7 | 12.27 | < 0.0001 |  |  |

**Table S5.** Path coefficients from two SEMs examining log-transformed synchrony (log φ) and community stability ($\log_{10} I_{c}$ ) as functions of diversity. Models were fit separately with richness and Shannon diversity as exogenous predictors. Entries are standardized coefficients (STD β) with *p*-values. Indirect effects are derived as Diversity → Synchrony × Synchrony → Stability.

| Response | Predictor | STD β | 95% CI | *p*-value | Interpretation |
| --- | --- | --- | --- | --- | --- |
| Model: Richness | | | | | |
| logφ | Richness | –0.178 | [-0.309, -0.047] | 0.008 | Richness weakly reduces synchrony |
| $\mathbf{log}_{\boldsymbol{10}} \boldsymbol{I}_{\boldsymbol{c}}$ | log φ | –0.911 | [-0.962, -0.860] | <0.001 | Synchrony strongly reduces stability |
| $\mathbf{log}_{\boldsymbol{10}} \boldsymbol{I}_{\boldsymbol{c}}$ | Richness | +0.033 | [-0.024, 0.090] | 0.31 | No direct stabilizing effect |
| Indirect | — | +0.162 |  | — | Modest positive indirect effect |
| Model: Shannon | | | | | |
| log φ | Shannon | –0.701 | [-0.789, -0.613] | <0.001 | Shannon strongly reduces synchrony |
| $\mathbf{log}_{\boldsymbol{10}} \boldsymbol{I}_{\boldsymbol{c}}$ | log φ | –0.851 | [-0.914, -0.788] | <0.001 | Synchrony strongly reduces stability |
| $\mathbf{log}_{\boldsymbol{10}} \boldsymbol{I}_{\boldsymbol{c}}$ | Shannon | +0.117 | [0.046, 0.188] | 0.0014 | Moderate direct stabilizing effect |
| Indirect | — | +0.597 |  | — | Large positive indirect effect |

**Table S6.** Linear mixed-effects model testing the effect of dissimilarity (Bray–Curtis dissimilarity between consecutive years) on community stability (log₁₀(I_C_)). The model includes season as a fixed effect (reference = Fall) and bay (major area) as a random intercept. Back-transformed estimates (10^Estimate) indicate multiplicative effects on I_C_ relative to the reference. Random intercept variance among bays = 0.024 (SD = 0.156); residual variance = 0.090 (SD = 0.300).

| Predictor | Estimate (log₁₀) | Std. Error | df | t | *p-value* | Back-transformed effect on $\boldsymbol{I}_{\boldsymbol{c}}$ |
| --- | --- | --- | --- | --- | --- | --- |
| Intercept (Fall) | 1.863 | 0.114 | 91.2 | 16.30 | < 0.001 | 10^1.863^ ≈ 72.96 (baseline) |
| Dissimilarity (β) | –2.470 | 0.193 | 325.9 | –12.82 | < 0.001 | 10^(–2.470)^ ≈ 0.0034 (–99.7%) |
| Spring | –0.199 | 0.047 | 324.0 | –4.25 | < 0.001 | 10^(–0.199)^ ≈ 0.63 (–37%) |
| Summer | –0.132 | 0.046 | 324.1 | –2.85 | 0.0046 | 10^(–0.132)^ ≈ 0.74 (–26%) |
| Winter | –0.038 | 0.047 | 324.1 | –0.81 | 0.42 | 10^(–0.038)^ ≈ 0.92 (–8%) |
