## Appendix S2 for "Evenness and Taylor’s law scaling shape biodiversity-stability relationships in subtropical estuarine communities"

**Journal:** Ecology

### Appendix S2. Supplementary Figures

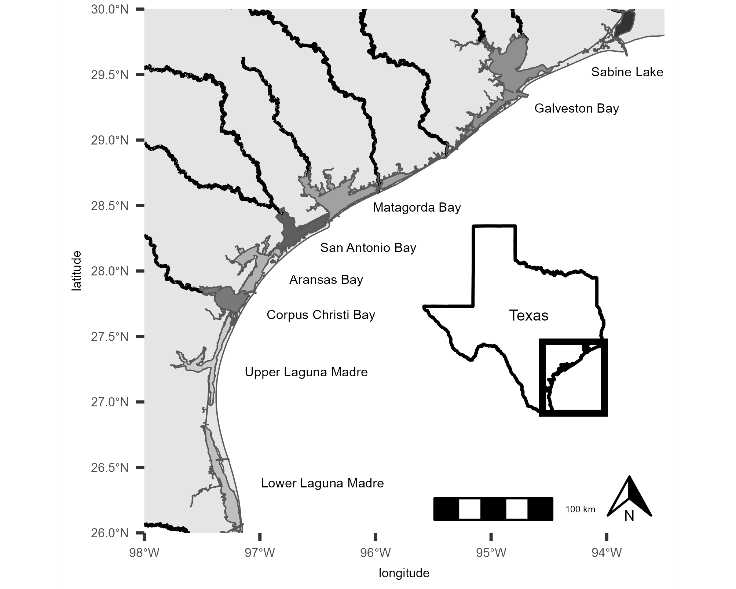

**Figure S1.** Study area along the Texas Gulf Coast. Shaded areas delineate the major estuaries analyzed: Sabine Lake; Galveston, Matagorda, San Antonio, Aransas, and Corpus Christi bays, Upper Laguna Madre, and Lower Laguna Madre.

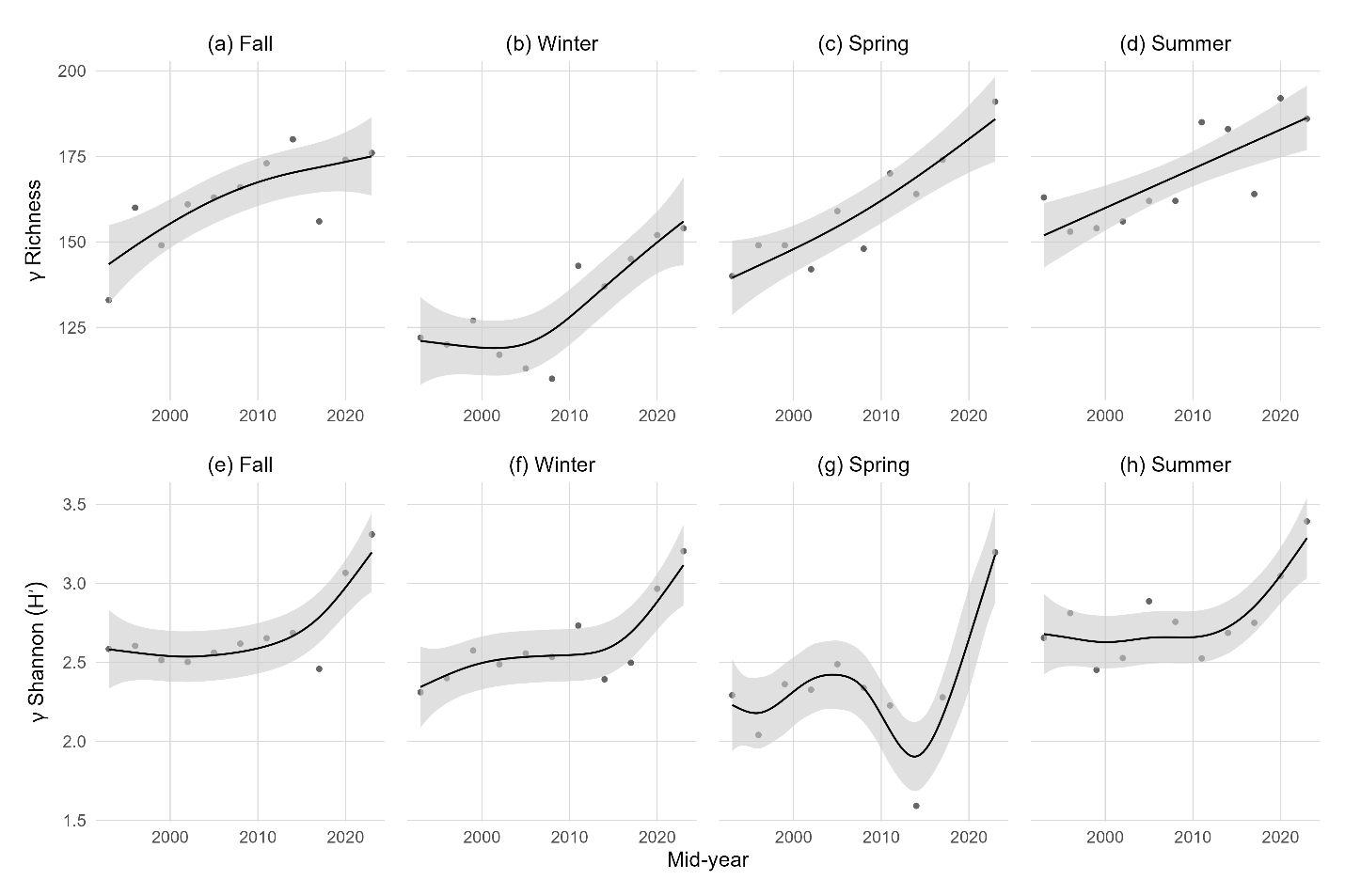

**Figure S2.** Seasonal generalized additive model (GAM) trends in regional γ-diversity. Points show observed regional γ-diversity pooled across bays within 3-year windows; the x-axis marks the mid-year of each window. Top row: γ richness (Hill number q = 0; panels (a)–(d) = Fall, Winter, Spring, Summer). Bottom row: γ Shannon diversity (Hill number q = 1; panels (e)–(h) = Fall, Winter, Spring, Summer). Black curves are GAM fits of γ-diversity vs. time (spline basis k = 6); gray ribbons show ≈95% confidence intervals (±2 SE). Separate y-axis scales are used for richness and Shannon.

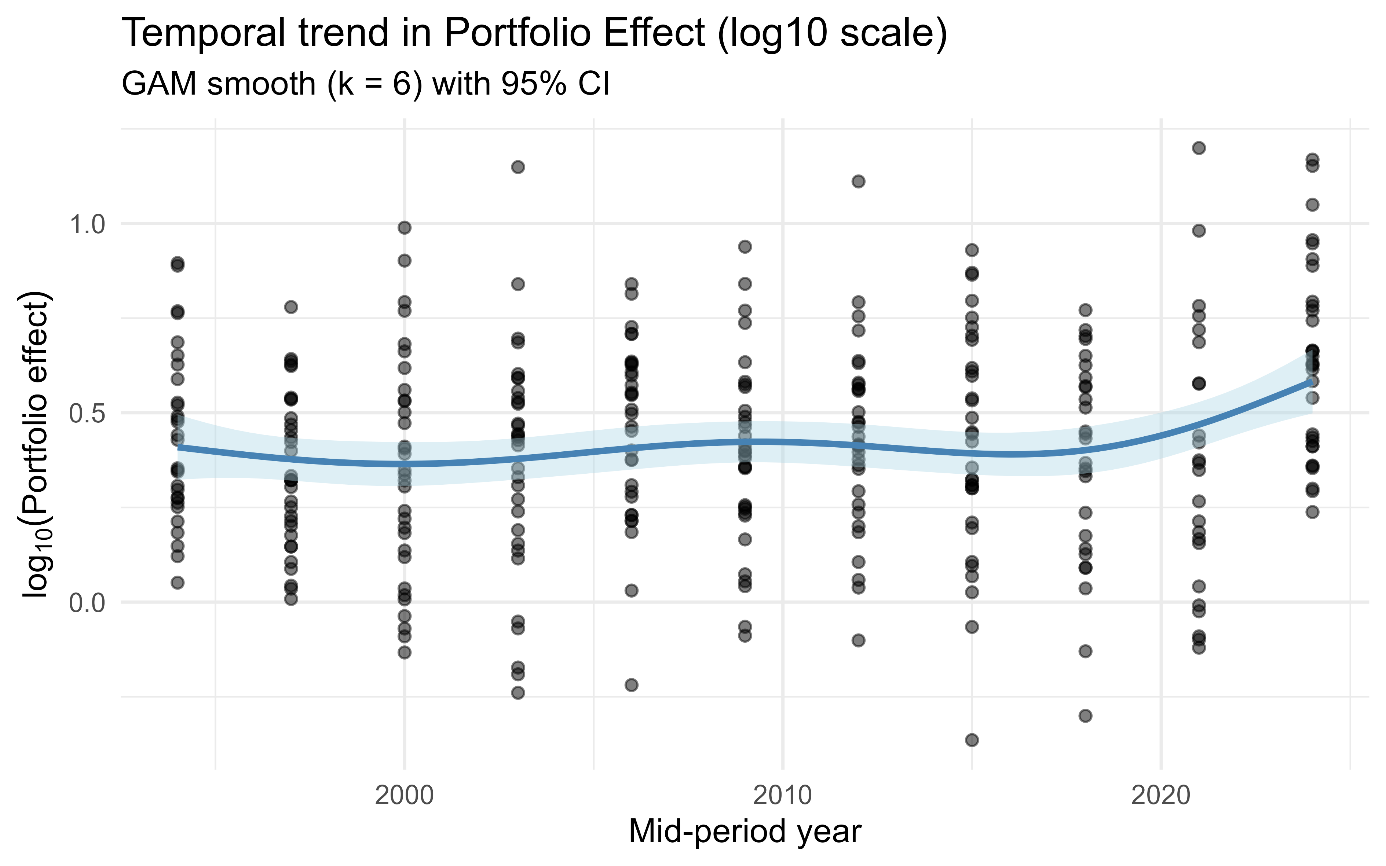

**Figure S3.** Temporal trend in the portfolio effect (log₁₀ scale) across bays and seasons. Points represent observed log₁₀(Portfolio) values, defined as the ratio of community-level invariability to the mean population invariability. The solid line shows the fitted generalized additive model (GAM) with a smooth function of mid-period year (k = 6), and the shaded region denotes the 95% confidence interval. The GAM revealed significant nonlinear temporal structure (smooth term: F₃.₉₉, ₄.₅₆ = 4.82, p < 0.001), explaining 27.4% of deviance (adjusted R² = 0.242; n = 336). Values > 1 indicate persistent statistical averaging via the portfolio effect.

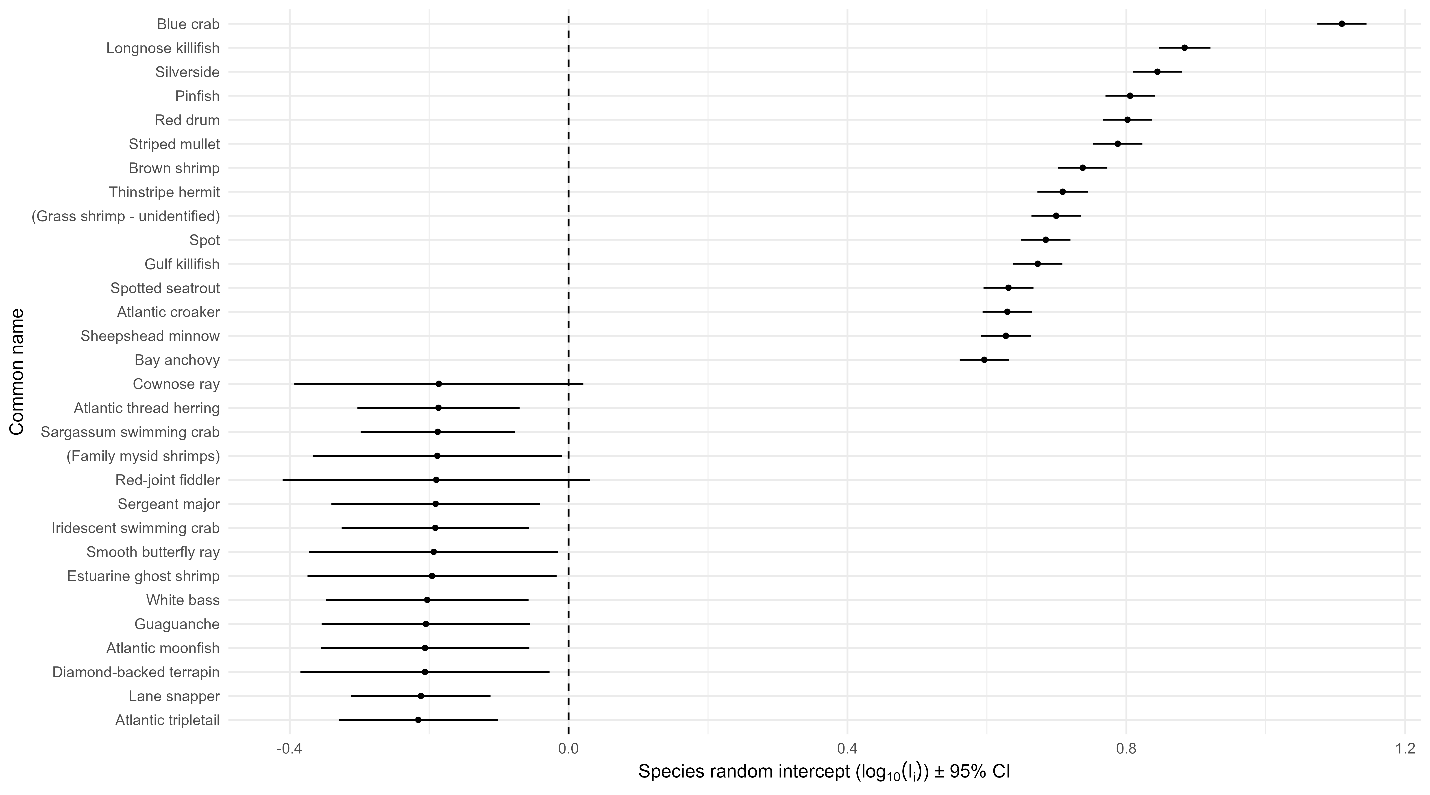

**Figure S4.** Top 15 species with the highest and lowest baseline contributions to community stability. Points show species-specific random intercepts from linear mixed-effects models of invariability ($\log_{10} I_{i}$) with season as a fixed effect and bay and species as random intercepts, with horizontal lines representing 95% confidence intervals. Positive values indicate species that contribute to higher baseline stability, whereas negative values indicate species associated with greater variability.
